# TCF-1 in CD4 T cells regulates GVHD severity and persistence

**DOI:** 10.1101/2021.03.22.436492

**Authors:** Rebecca Harris, Mahinbanu Mammadli, Adriana May, Qi Yang, Ivan Ting Hin Fung, Jyoti Misra Sen, Mobin Karimi

## Abstract

Graft-versus-host disease (GVHD) is a leading cause of mortality following allogeneic hematopoietic stem cell transplantation (allo-HSCT). Mature donor T cells in the graft mediate graft-versus-leukemia (GVL) responses against residual tumor cells, which may persist after pre-transplant conditioning regimens. Importantly, the same mature T cells also mediate GVHD. The transcription factor T Cell Factor-1 (TCF-1) is critical for T cell development in the thymus. Using a unique mouse model of allo-HSCT leading to GVHD, we investigated the role of TCF-1 in alloactivated T cell functioning and in GVHD. Here, we report that loss of TCF-1 in mature CD4 T cells reduces GVHD severity and persistence, improving survival of recipient mice. This was due to reduced proliferation, survival, and cytokine production of T cells, as well as increased exhaustion. Gene pathways involved in cytokine response, immune signaling, chemokine signaling, cell cycle, and T cell differentiation were altered by loss of TCF-1 in donor cells. Our companion paper shows that regulation of alloactivated CD4 T cells by TCF-1 differs from regulation of CD8 T cells, suggesting that TCF-1 plays a unique role in each subset. Therefore, targeting of TCF-1 or downstream signaling pathways may be an effective strategy for reducing GVHD following allo-HSCT.

## Introduction

Patients with hematological malignancies often undergo allogeneic hematopoietic stem cell transplantation (allo-HSCT), a curative therapy (1). This involves donor stem cells being transplanted, along with mature donor T cells (2). These mature T cells in the graft help to clear residual malignant cells and prevent relapse, an outcome called the graft-versus-leukemia (GVL) effect. However, 30-70% of these patients also develop graft-versus-host disease (GVHD), a life-threatening complication mediated by these same donor T cells (3). Alloactivation of the T cells by non-self HLA leads to T cell proliferation, migration, and production of cytokines, resulting in damage to healthy tissues (4). GVHD remains a major clinical barrier to widespread use of allo-HSCT, and depletion of T cells or immunosuppressive therapies are sub-optimal, leaving the patient susceptible to infection or relapse (5). Our research seeks ways to modulate mature T cell signaling during alloactivation to separate the linked processes of GVHD and GVL. To reach this goal, better understanding is needed on how alloactivated T cells are regulated.

T Cell Factor-1 (TCF-1) is a T-cell transcription factor which is critical for T cell development (6, 7). TCF-1 is also known to be involved in T cell proliferation, recall, and memory responses during viral infection (8). However, to our knowledge, TCF-1 has not been studied in the context of alloactivation, which is a different process from canonical activation – resulting from recognition of non-self HLA rather than of pathogens (9). TCF-1 has also been reported to affect the transcription factor Eomesodermin (Eomes) and possibly T-box transcription factor 21 (T-bet) (10, 11). Eomes and T-bet affect both GVHD and GVL, but it remains unknown whether TCF-1 may play a role upstream of these factors in separating GVHD from GVL (12–15).

Global deletion of TCF-1 significantly reduces the number of thymocytes, as well as the number of mature T cells exiting the thymus, because TCF-1 is critical for the double-negative stages of development (16–18). To overcome this problem, we obtained mice that have a T cell-specific deletion of TCF-1 using CD4cre (Tcf7 flox/flox x CD4cre) (19, 20). These mice are able to produce mature T cells because they retain TCF-1 expression until the double-positive phase of development in the thymus(20). At this point, all T cells will have TCF-1 expression silenced by CD4cre (because all T cells express CD4 at this point, even CD8 T cells)(20). Therefore, this unique mouse strain allows us to study mature T cells with TCF-1 deficiency.

Using a mouse model of GVHD and GVL following allo-HSCT, we found that loss of TCF-1 in CD4 T cells reduces both the severity and the persistence of GVHD, leading to improved survival of recipient mice following transplant. This disease reduction was driven by alterations to T cell proliferation, cytokine production, survival, and exhaustion, as well as changes in gene expression. Expression of chemokine receptors on donor T cells was enhanced by loss of TCF-1. We also showed that loss of TCF-1 in CD4 T cells induces T cell-intrinsic phenotypic changes to memory identity, including increased CD4 naive cells and decreased activating/transitioning CD4 T cells. TCF-1 also affects expression of genes in the cytokine response, cell cycle control, T cell differentiation, and immune signaling pathways, among others. Therefore, TCF-1 controls mature CD4 T cell phenotype, functions, and gene expression during alloactivation, a novel finding.

Here, we provide evidence that TCF-1 regulates mature alloactivated T cell function and genetic programs, thereby controlling GVHD severity and persistence. The major alloactivated T cell functions – cytokine production, migration, and proliferation – were altered by loss of TCF-1, as were cell survival and gene expression. Our companion manuscript shows that the control of CD4 T cells by TCF-1 is distinct from control of CD8 T cells. Therefore, TCF-1 acts as a regulatory transcription factor in mature alloactivated CD4 T cells, and modulation of TCF-1 and downstream signaling may be a successful option for reducing GVHD to improve patient outcomes following allo-HSCT.

## Results

### Loss of TCF-1 in donor CD4 T cells reduces severity and persistence over time of GVHD symptoms

Previous research on TCF-1 has focused on T cell development and canonical activation (21, 22). These studies primarily used a global TCF-1 knockout, resulting in limited production of mature T cells (21, 22). We sought to examine the role of TCF-1 in mature T cells, so to overcome this limitation we employed mice with T-cell-specific deletion of TCF-1 using CD4cre (20). This mouse strain has TCF-1 deleted in all CD4 and CD8 T cells at the DP phase, allowing production of mature T cells with TCF-1 deletion. To investigate whether TCF-1 in mature CD4 T cells contributes to GVHD after allo-HSCT, we employed a murine model of MHC-mismatched allotransplantation. Donor CD4 T cells from C57Bl/6 background wild-type (WT), CD4cre control, or TCF-1-deficient (TCF cKO) mice were injected along with T cell depleted BALB/c bone marrow into lethally irradiated BALB/c recipients as described in the methods (15, 23). The MHC haplotype mismatch (H2Kb in donors, H2Kd in recipients) results in alloactivation of the donor T cells, leading to GVHD (15). Recipient mice were then weighed and given a GVHD score (Fig.1A) to identify the severity of GVHD for up to 60 days post-transplant. Recipient mice were scored based on weight loss, fur texture, posture, activity, skin condition, and diarrhea, as previously described(15, 24).

**Figure 1:**
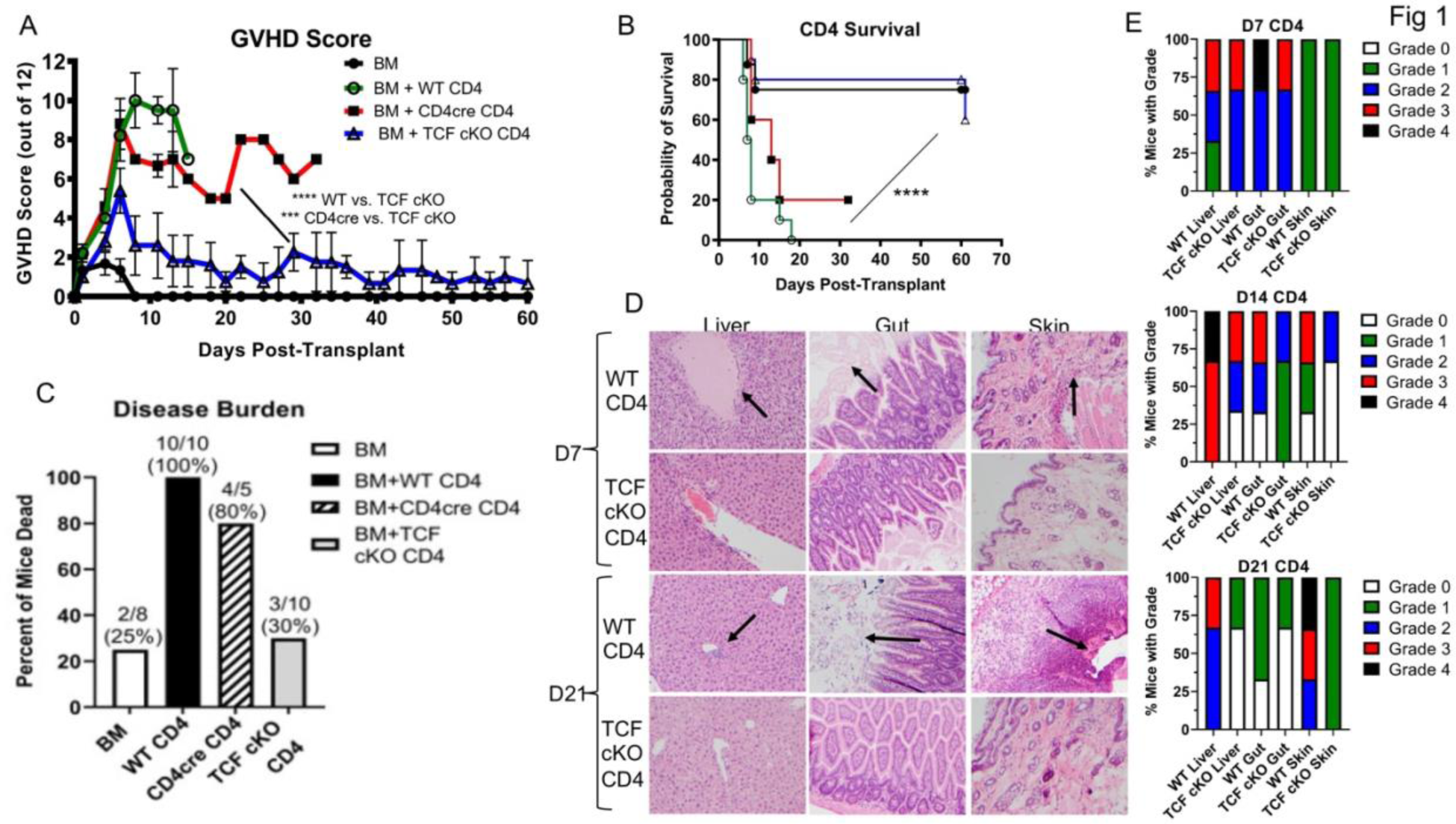
Loss of TCF-1 in donor CD4 T cells reduces severity and persistence over time of GVHD symptoms. Recipient BALB/c mice (MHC haplotype D) were lethally irradiated and allotransplanted with 10×10^6^ BALB/c bone marrow (BM) cells and 1×10^6^ CD4 T cells from WT, CD4cre, or TCF cKO donor mice (C57Bl6 background, MHC haplotype B). The mice were weighed and given a GVHD clinical score three times per week for 60 days. Score was determined by combined scores for fur texture, activity level, posture, skin integrity, weight loss, and diarrhea. **(A)** Clinical scores for mice over 60 days. Mean and SD plotted, analyzed by two-way ANOVA. **(B)** Survival of mice in each group over the 60 day experiment, analyzed with Kaplan-Meier survival statistics. **(C)** Disease burden as determined by percent of mice in each group that died from GVHD by the end of the experiment. Number of mice (dead and total) per group as well as percent dead are noted above each bar. **(D)** Organs (skin, liver, and small intestine AKA “gut”) from recipient mice taken at day 7, day 14, and day 21 post-transplant, sectioned, and stained with H&E. A pathologist (A.M.) who was blinded to study conditions and sample groups was given deidentified slides, and gave each tissue section a GVHD grade based on damage and other indicators (detailed further in Methods). Shown here are representative sections from D7 and D21 for each organ. Arrows indicate areas of damage or lymphocytic infiltration. **(E)** Histology GVHD grades were graphed as percentage of mice in each group with that grade. Grades 0-2 are considered mild GVHD and grades 3-4 are considered severe GVHD. Chi-square analyses were used to determine differences between grade frequencies. **** means p-value ≤ 0.0001. N=3-5 per group for **A-B** with one representative experiment shown, N=5-10 per group in C with two experiments combined, N=3 per group for D-E, one representative photo shown and summary data in E.

Mice receiving CD4 T cells from WT or CD4cre donors experienced a rapid increase in GVHD symptoms, peaking at a very high score, indicating very severe disease (25). CD4 T cells are known to cause very severe GVHD symptoms, so this finding was expected (25). Over time, these recipient mice continued to show severe symptoms, with consistent scores until death from disease (Fig.1A). Survival was also poor, with most mice in this group dying prior to day 10 (Fig.1B). These mice lost weight due to disease, and died before they were able to regain much weight (Supp. Fig.1A). The disease burden, as determined by total number of mice per group that died by the end of the experiment, was very high for WT and CD4cre-transplanted mice (Fig.1C). In contrast, mice receiving CD4 T cells from TCF cKO mice showed reduced GVHD scores (Fig.1A), better survival (Fig.1B), and weight gain following initial weight loss (Supp. Fig.1A). Disease burden was reduced in this group (Fig.1C), with most mice in this group remaining alive at day 60 post-transplant. Interestingly, mice in the TCF cKO donor group still showed a peak in GVHD score early on, as with the WT group, but this peak was at a much lower score. Additionally, this peak score did not persist over time, as the scores for these mice quickly reduced to the level seen in bone marrow-only controls (Fig.1A). In addition, this low score remained low for an extended period of time, suggesting that disease had resolved rather than been delayed (Fig.1A). These data indicated that GVHD symptoms were not only less severe in these mice, but also less persistent over time.

To examine pathological damage to target organs – including the liver, guts, and skin – tissues from these organs were collected for histology at three time points. At day 7, day 14, and day 21 post-transplant, organs were taken for histology, H&E stained, and scored for GVHD pathology. At day 7, little to no difference was observed in tissue damage or GVHD grade between WT and TCF cKO transplanted mice (Fig.1D-E), (Supp. Fig.1B), in agreement with the GVHD score data, where major differences occurred after day 7. By day 21, differences in GVHD grade were observed between WT and TCF cKO recipients, with lymphocyte infiltration visible in liver, guts, and skin of WT but not TCF cCKO-transplanted mice (Fig.1D-E), (Supp. Fig.1C). In addition, tissue damage in guts and skin are clearly visible for WT but not TCF cKO-transplanted mice (Fig.1D). Splenic structural disorganization is also visible in spleen of WT but not TCF cKO-transplanted mice (Supp. Fig.1C). By day 14, these changes are already visible as a trend in GVHD grade, indicating a progressive resolution of GVHD symptoms over time in TCF cKO-transplanted mice (Supp. Fig.1B). By day 21, the percentage of mice experiencing severe GVHD (grades 3-4, versus no or mild to moderate GVHD at grades 0-2.(15, 26)(27) was trending towards increased for WT-transplanted mice, compared to TCF cKO-transplanted mice, and this trend can be seen as early as day 14 (Fig.1E). At day 7, there is no apparent difference between the distribution of disease grades (Fig.1E). For all organs, the histology grade tends to decrease over time for TCF cKO-transplanted mice, but rise or remain consistent for WT-transplanted mice (Supp. Fig.1B), further supporting the idea that disease resolves over time and does not persist when donor cells are TCF-1-deficient. Together, these results indicate that TCF-1 normally contributes to and is indispensable for GVHD damage by T cells, and loss of TCF-1 reduced severity and persistence of GVHD.

### Loss of TCF-1 drives changes to mature CD4 T cell memory identity which are primarily cell-intrinsic

TCF-1-deficiency in CD4 T cells alters GVHD outcomes, suggesting that T cell phenotypes or function are affected. To examine the phenotype of mature CD4 T cells when TCF-1 is lost, we performed flow cytometry phenotyping on naive CD4 T cells from WT and TCF cKO mice (Fig.2A-B), (Supp. Fig.2A-C). We began by looking at expression of the transcription factors Eomesodermin (Eomes) and T-box transcription factor 21 (T-bet), which are downstream of TCF-1. Previous reports suggested that TCF-1 normally activates Eomes, and may activate or not impact T-bet (28). However, we found that loss of TCF-1 in CD4 T cells did not affect expression of Eomes and T-bet (Supp. Fig.2A-B). In addition, expression of CD122 was not impacted by loss of TCF-1 (Supp. Fig.2C). Interestingly, expression of CD44 was polarized more potently in TCF cKO T cells (Fig.2A-B). Using CD62L and CD44, we identified four major memory subsets: central memory (CDD44+ CD62L+), effector memory (CD44+ CD62L-), naive (CD44-), and activating/transitioning (CD44 mid). Comparing WT and TCF cKO mice, we found that loss of TCF-1 results in an increase in CD4+ naive T cells and a decrease in CD4+ transitioning/activating cells compared to WT mice (Fig.2A). These results suggest that loss of TCF-1 results in a more naive phenotype in CD4+ T cells. Given the increased ability of naive cells to cause severe GVHD, this would suggest that loss of TCF-1 would increase GVHD severity. Thus, some other T cell functions must be impacted to explain the reduction observed.

**Figure 2:**
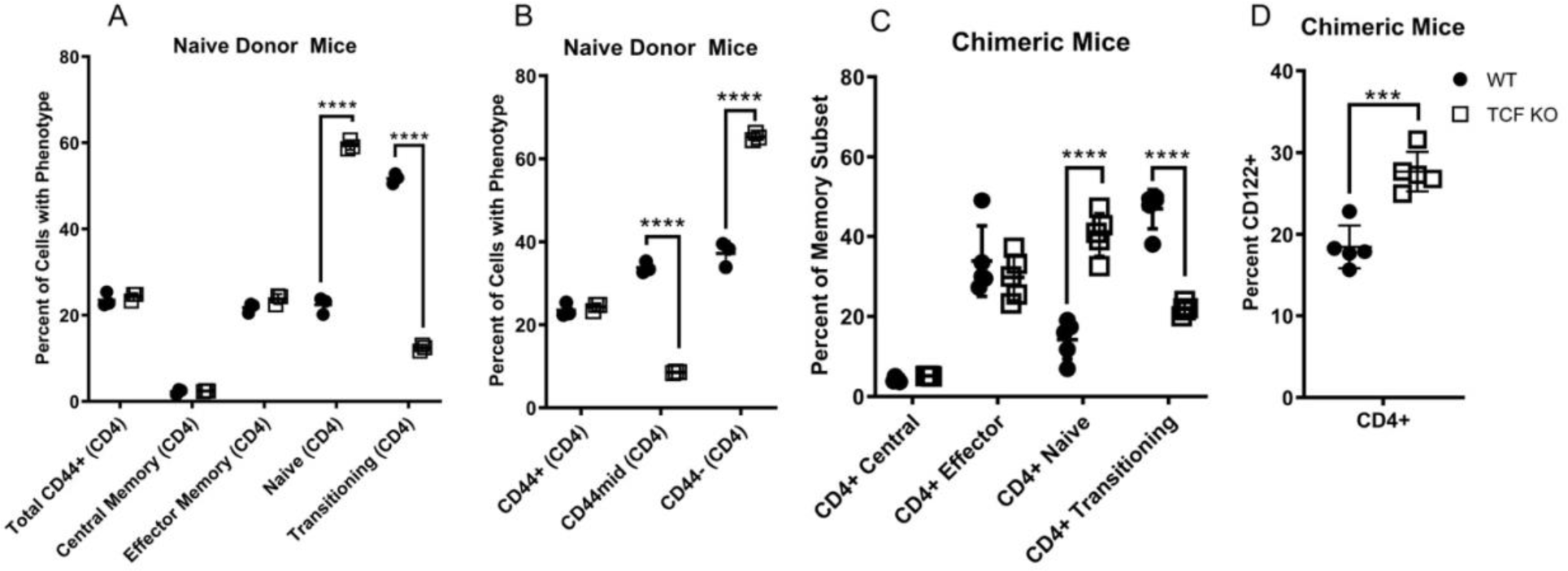
Loss of TCF-1 drives changes to mature CD4 T cell memory identity which are primarily cell-intrinsic. (A-B) Naive WT or TCF cKO donor mice were euthanized, splenocytes were obtained and stained for flow cytometry, and run on a BD LSRFortessa flow cytometer. **(A)** Percent of CD4 T cells expressing naive, transitioning/activating, central memory, or effector memory phenotypes. (B) Percent of CD4 T cells expressing CD44. **(C-D)** Thy1.1 mice were lethally irradiated and reconstituted with a 1:4 (WT:TCF cKO) mixture of bone marrow cells. At 9 weeks, blood was checked by flow cytometry to ensure reconstitution, and at 10 weeks, flow cytometry phenotyping was performed. WT donor cells were identified by CD45.1, while TCF cKO donor cells were identified by CD45.2 **(C)** Percent of chimeric CD4 T cells expressing naive, transitioning/activating, central memory, or effector memory phenotypes. **(D)** Percent of chimeric CD4 T cells expressing CD122. All data are shown as individual points with mean and SD, all data were analyzed with Student’s t-test, one-way ANOVA, or two-way ANOVA (depending on data groups), **** means p-value ≤ 0.0001, and *** means p-value ≤ 0.001. For A-B, N=3 per group with one representative experiment shown, for C-D, n=5 per group, one experiment shown (done once).

To investigate whether these changes were cell-extrinsic or -intrinsic, we utilized a chimeric bone marrow mouse model, as described in the methods. Briefly, Thy1.1 mice were lethally irradiated and reconstituted with bone marrow from WT and TCF cKO donors, mixed at a 4:1 (TCF:WT) ratio, for a total of 50×10^6^ BM cells. Due to our initial observations that TCF cKO T cells did not proliferate well in culture, we used a 4:1 ratio to ensure survival of enough TCF cKO bone marrow cells to perform downstream analyses. At 10 weeks post-transplant, splenocytes were obtained from the recipient mice and analyzed by flow cytometry, with donor cells identified by H2Kb expression. The increase in naive CD4+ T cells, as well as the reduction in activating cells, was found to be cell-intrinsic in the TCF cKO T cells, because the differences were maintained despite development in a chimeric mouse (Fig.2C). Thus, the memory identity changes induced by loss of TCF-1 are cell-intrinsic. Of note, CD122 expression was increased among TCF cKO donor cells from chimeric mice (Fig.2D), suggesting that these cells are capable of being activated. Again, no differences were seen in Eomes and T-bet expression (Supp. Fig. 2D-E).

### Loss of TCF-1 induces an apparent migration defect and changes to chemokine/chemokine receptor expression in mature CD4 T cells during alloactivation

Our data show that GVHD is disrupted when TCF-1-deficient CD4 T cells are used for allotransplantation. We next sought to determine which of the major alloactivated T cell functions - migration, proliferation, and cytokine production – were altered to provide the optimal clinical phenotype. First, we examined CD4 T cell migration by transplanting lethally irradiated BALB/c mice with WT and TCF cKO total CD3+ T cells mixed in a 1:1 ratio (1×10^6^ total cells), along with T cell-depleted BALB/c bone marrow (BM, 10×10^6^ cells). The donor cells were checked prior to transplant for CD45.1/CD45.2 (WT and TCF cKO congenic markers, respectively) (Fig.3A), (Supp. Fig.3). CD4 and CD8 T cells were first mixed 1:1 for each strain, then a 1:1 mixture of the two strains was prepared. WT CD45.1 and WT CD45.2 cells were mixed in the same way as a control. After 7 days, the spleen, lymph nodes, small intestine (SI or “gut”), and liver were removed from recipients and lymphocytes were obtained (15).

**Figure 3:**
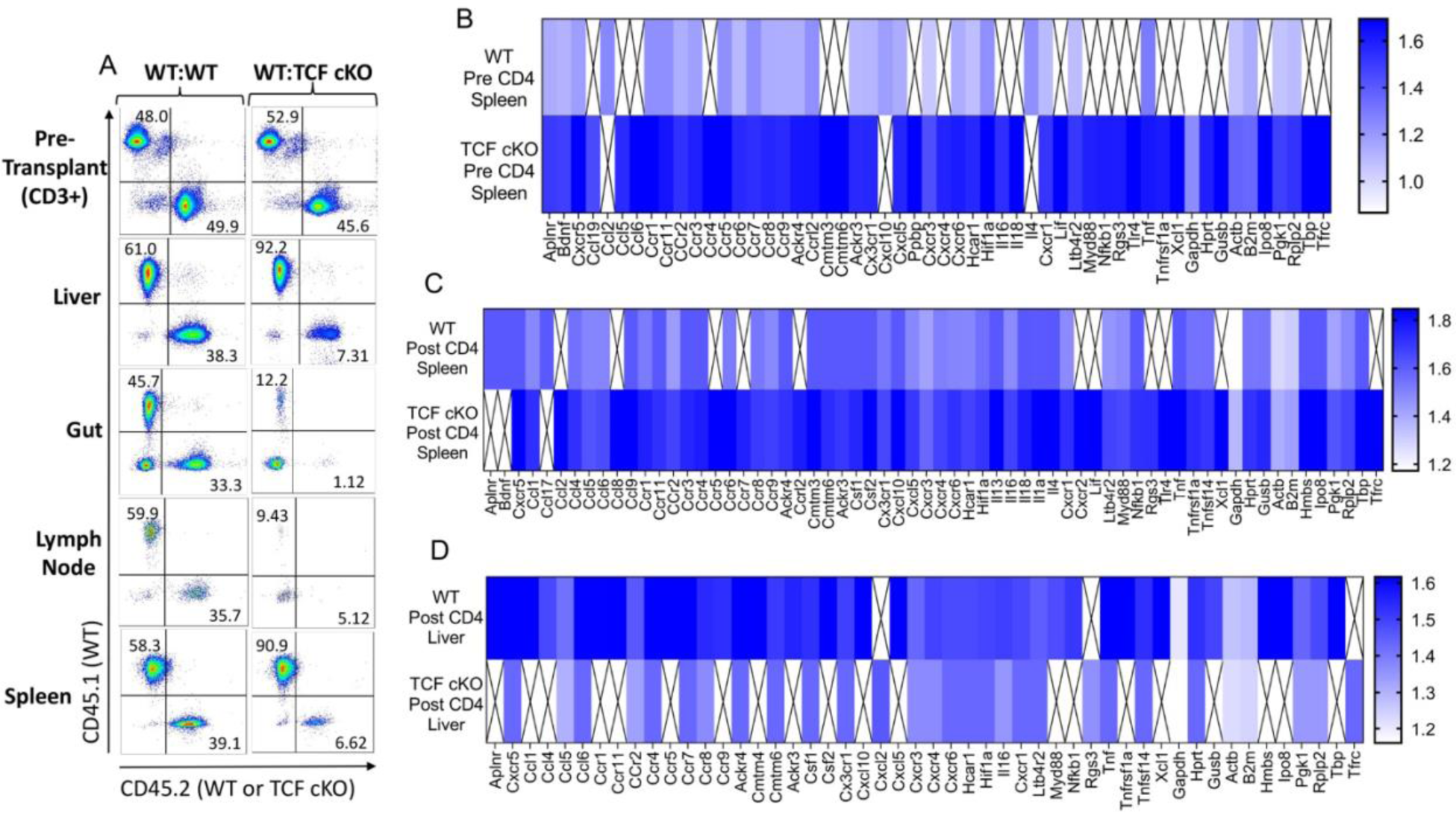
Loss of TCF-1 induces an apparent migration defect and changes to chemokine/chemokine receptor expression in mature CD4 T cells during alloactivation. Recipient BALB/c mice were lethally irradiated and allotransplanted with 10×10^6^ BALB/c bone marrow (BM) cells and 1×10^6^ CD3 T cells. The T cells were a 1:1 mixture of CD3 T cells from WT (CD45.1) and TCF cKO (CD45.2) donor mice, or WT (CD45.1) and WT (CD45.2) control mice. **(A)** Prior to transplant, the cell mixture was checked to ensure a 1:1 ratio of CD4/CD8 T cells, and a 1:1 ratio of each donor strain (using CD45.1 and CD45.2). At day 7 post-transplant, the spleen, liver, small intestine (“gut”), and lymph nodes were taken from euthanized recipient mice and processed to obtain lymphocytes. The cells were then stained for H2Kb (donor cells), H2KD (recipient cells), CD3, CD4, CD8, CD45.1, and CD45.2. The pre-transplant flow plots are shown on top, and one representative post-transplant flow plot for WT:WT or WT:TCF cKO mice is shown per organ below. **(B-D)** To perform qPCR, BALB/c mice were lethally irradiated and allotransplanted with 1×10^6^ CD3 donor cells and BALB/c BM, as above. Donor CD4 T cells from WT and TCF cKO mice were FACS sorted both pre-transplant and 7 days post-transplant, from spleen (both pre and d7) and liver (d7 only). The cells were sorted in Trizol, and RNA was extracted using chloroform and converted into cDNA using a synthesis kit (more details in Methods). cDNA was added to premade mouse chemokine/chemokine receptor primer plates (ThermoFisher), and run on a QuantStudio 3 thermocycler. Results are shown as a heatmap of fold change per gene, compared to 18S reference gene for each plate. Boxes with an “X” represent signals too low to detect or otherwise unreadable due to technical error. Heatmaps compare WT versus TCF cKO CD4 T cells in **(B)** pre-transplant spleen, **(C)** post-transplant spleen, and **(D)** post-transplant liver. N=3 per group for A with one representative experiment shown, N=5 mice into one sample per condition for **B-D**, summary data shown.

For CD4+ T cells, TCF cKO cells were present at lower frequencies in the liver and SI, indicating a potential migration defect (Fig.3A), (Supp. Fig. 3). Based on these data, the frequency of TCF cKO CD4 T cells would be expected to be higher than WT in the spleen and lymph nodes, where the cells would be trapped. However, the frequency of TCF cKO CD4 T cells was still lower than WT in these organs (Fig.3A), (Supp. Fig.3). This suggested that in CD4 TCF-1-deficient T cells, a migration defect may exist, but this could also be the result of reduced survival or a potential defect in proliferation.

To determine whether specific changes to chemokines or chemokine receptors were responsible for the migration defect, and to examine whether a true migration defect existed, we sorted back donor CD4 T cells from allotransplanted BALB/c mice (using H2Kb to identify donor cells) and performed qPCR using a 96-well mouse chemokine/chemokine receptor array plate (ThermoFisher). We found that the expression of chemokines and chemokine receptors was upregulated following alloactivation, as expected. However, expression of these markers was consistently higher in TCF cKO CD4 T cells from spleen, both pre- and post-transplant (Fig.3B-C), while these markers were downregulated in TCF cKO CD4 T cells from post-transplant liver (Fig.3D). Therefore, CD4 T cells from TCF cKO mice should be capable of migration at normal levels, and the observed difference in migration is likely due to changes in proliferation or cell survival. However, it is possible that these cells may have a mild migration defect, which could become more apparent later in the disease course.

### TCF-1 controls proliferation, apoptosis, and exhaustion of mature alloactivated CD4 T cells

The donor T cells responsible for GVHD must proliferate in the secondary lymphoid organs and in the GVHD target organs in order for the response to continue (29). Effector T cells - which drive GVHD - have a short lifespan, so they must be continually replaced during an alloresponse. To investigate whether loss of TCF-1 affects the proliferation of mature alloactivated T cells, we transplanted lethally irradiated BALB/c mice with 1×10^6^ WT or TCF cKO CD3 T cells and 10×10^6^ BALB/c BM, as described above. The recipient mice were injected with EdU in PBS at 25mg/kg on day 5-6. At day 7, recipient mice were euthanized, lymphocytes were obtained from the spleen and liver, and donor cells (identified by H2Kb, CD3, and CD4) were tested for the presence of EdU with a click chemistry flow cytometry kit. A trend of increased proliferation in the spleen and significantly increased proliferation in the liver by TCF cKO donor CD4 T cells was observed (Fig.4A-B). Therefore, proliferation at day 7 is increased in TCF cKO CD4 T cells. We also used expression of Ki67 in a similar experiment to examine the activation or proliferation of these donor cells, and found that activation/proliferation of T cells was trending towards increased for TCF donor cKO CD4 T cells in the spleen, but not in the liver (Fig.4C-D). Therefore, proliferation in T cells early after alloactivation is increased by loss of TCF-1. However, given that symptoms of GVHD resolve after day 7 in TCF cKO-transplanted recipients, we hypothesize that proliferation of donor cells also decreases below WT levels after this point.

**Figure 4:**
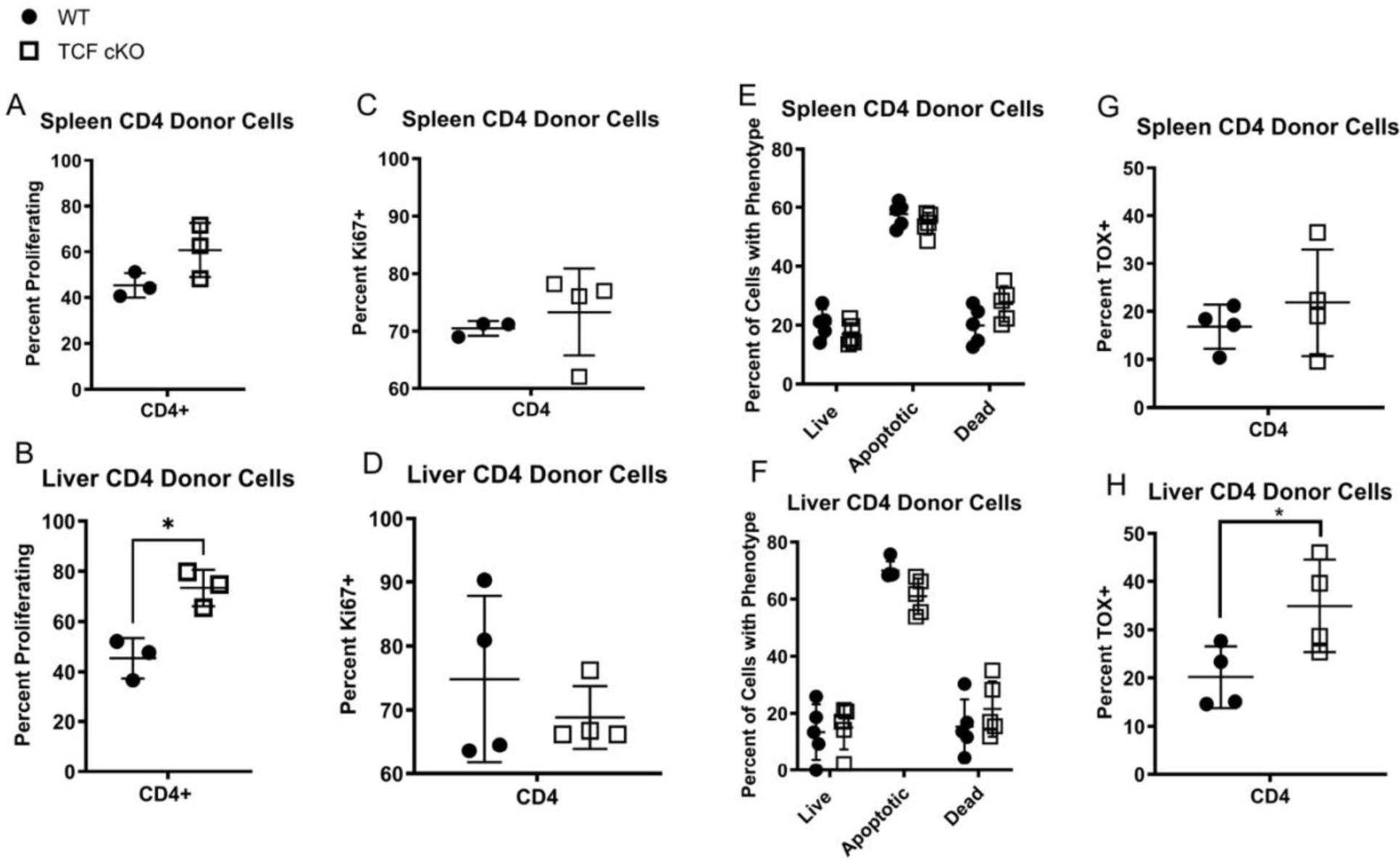
TCF-1 controls proliferation, apoptosis, and exhaustion of mature alloactivated CD4 T cells. (A-B) Recipient BALB/c mice were allotransplanted with BALB/c BM and 1×10^6^ WT or TCF cKO donor CD3 T cells, as before. On day 5 and 6, recipient mice were injected i.p. with 25mg/kg of EdU in PBS. At day 7, spleens and livers were removed from euthanized recipient mice, lymphocytes were isolated, and stained for EdU with a click chemistry kit. Cells were also stained for H2Kb, CD3, CD4, and CD8. Graphs show percent proliferating (by EdU+) in **(A)** spleen and **(B)** liver. **(C-D)** Recipient mice were allotransplanted as before, and at day 7 spleens and livers were obtained from euthanized recipients. Lymphocytes were isolated, and cells were stained for H2Kb, CD3, CD4, CD8, and Ki67 to identify proliferating/activated cells in **(C)** spleen and **(D)** liver. **(E-F)** Recipient mice were allotransplanted as before, and at day 7, spleens and livers were taken from euthanized recipients. Lymphocytes were isolated and stained for CD3, CD4, CD8, H2Kb, and with Annexin V-FITC and LIVE/DEAD Near IR. Cells were identified as live (Ann.V-IR-), apoptotic(Ann.V+IR-), or dead (Ann.V+IR+), in both **(E)** spleen and **(F)** liver. **(G-H)** Recipient mice were allotransplanted as before, and at day 7 spleens and livers were obtained from euthanized recipients. Cells were stained for H2Kb, CD3, CD4, and CD8, then fixed/permeabilized and stained for TOX to identify exhausted cells in **(G)** spleen and **(H)** liver. All data are shown as individual points with mean and SD, were analyzed with one-way or two-way ANOVA, and * means p-value ≤ 0.05. N=3-5 per group with one representative experiment shown.

To examine the role of TCF-1 in survival of mature T cells during alloactivation, we performed a death assay using flow cytometry. Recipient mice were allotransplanted in the same manner as for proliferation experiments, and at day 7, cells from the spleen and liver were analyzed by flow cytometry. Donor cells were detected with H2Kb, CD3, CD4, and CD8. Using Annexin V-FITC and LIVE/DEAD Near IR, we were able to separate three distinct populations - live cells, dead cells, and apoptotic cells (Fig.4E-F). When we compared WT and TCF cKO donor cells, we found that TCF cKO CD4 T cells were trending towards more frequently dead, and less frequently live or apoptotic, than WT donor cells (Fig.4E-F). In support of this, when we attempted to sort CD4 T cells from WT- and TCF cKO-transplanted mice at day 7, there were significantly fewer CD4 donor T cells in the spleen from TCF cKO donors than from WT donors (by number of cells/10,000 total cells) (Supp. Fig. 4A-B). In addition, we attempted to sort TCF cKO donor cells from recipient mice at day 14 post-transplant, and donor cells in the spleen were significantly reduced (with cells in liver trending to reduced) compared to WT and TCF cKO donor T cells from day 7 (Supp. Fig. 4A-B). Empirical observations from our work also suggested that TCF cKO T cells proliferate poorly in culture compared to WT T cells, even when activated. Therefore, it is likely that survival and/or proliferation are reduced further after day 7, giving rise to the progressive resolution in symptoms observed in TCF cKO-transplanted mice.

Finally, to determine whether exhaustion of the T cells occurred when TCF-1 was lost, we allotransplanted mice as described above, and assessed donor cells from the spleen and liver at day 7 post-transplant for the expression of TOX, a T-cell exhaustion marker. We found that donor CD4 T cells from TCF cKO mice were significantly more exhausted in the liver, and trending towards more exhausted in the spleen, than WT donor cells (Fig.4G-H). Together, these data suggest that CD4 donor T cells from TCF cKO mice have worse survival, increased exhaustion, and a potential progressive defect in proliferation over time compared to WT CD4 T cells.

### Loss of TCF-1 changes the cytokine profile of mature alloactivated CD4 T cells

Production of inflammatory cytokines, eventually culminating in a cytokine storm, in considered a hallmark of GVHD (29, 30). Cytokines and cytotoxic mediators are essential for T cells to maintain the GVL effect and kill tumor cells, yet they also lead to damage of healthy host tissues (30, 31). To examine cytokine production by TCF cKO CD4 T cells, we allotransplanted recipient mice as described above with 1.5×10^6^ CD3 donor T cells (WT or TCF cKO) and 10×10^6^ BALB/c BM. At day 7 post-transplant, splenocytes were obtained from recipient mice, and were restimulated to induce cytokine production. Cells were restimulated with Golgiplug along with PBS (negative control) or anti-CD3/anti-CD28 (restimulated) for 6 hours at 37C. After this incubation, the cells were stained for flow cytometry with antibodies against H2Kb, CD3, CD4, CD8, TNF-α, and IFN-γ. We found that production of TNF-α by donor cells trended toward decreasing when TCF-1 was lost (Fig.5A). On the other hand, production of IFN-γ did not appear to be affected by loss of TCF-1 (Fig.5B).

**Figure 5:**
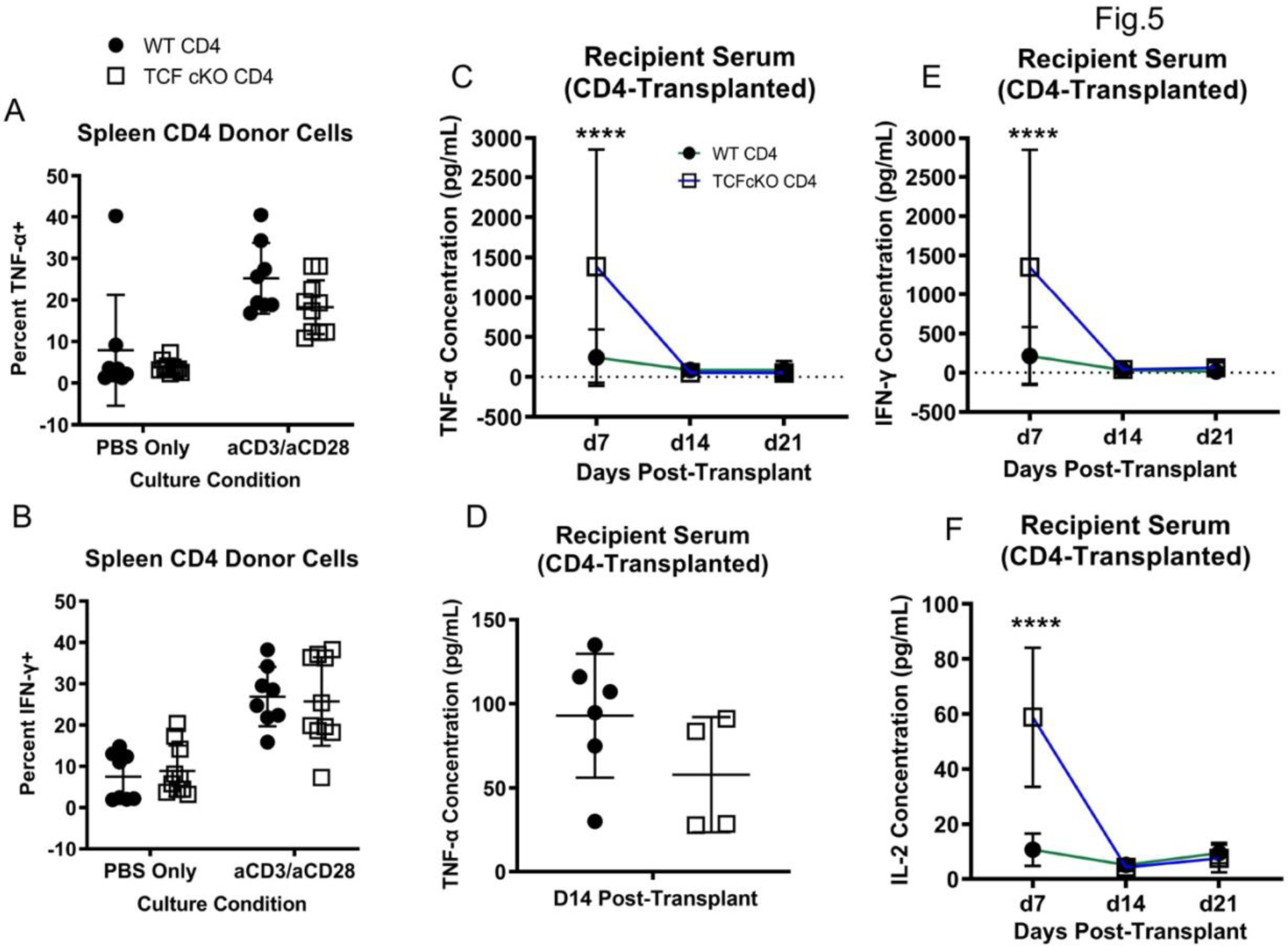
Loss of TCF-1 changes the cytokine profile of mature alloactivated CD4 T cells. (A-B) Recipient Balb/c mice were allotransplanted with Balb/c BM and 1.5×10^6^ WT or TCF cKO donor CD3 T cells, as before. At day 7, splenocytes were obtained and restimulated by 6 hours of culture with anti-CD3/anti-CD28 or PBS (control), along with GolgiPlug. Production of **(A)** TNF-α and **(B)** IFN-γ was measured by flow cytometry, using percent cytokine-positive cells. Donor cells were stained for H2Kb, CD3, CD4, and CD8, then fixed/permeabilized and stained with anti-IFN-γ and anti-TNF-α. **(C-F)** Serum was obtained from cardiac blood of euthanized recipient mice at day 7, day 14, and day 21 post-transplant, and was tested using a LEGENDplex multiplex ELISA kit. **(C)** Serum concentration (pg/mL) of TNF-α over time for WT versus TCF cKO-transplanted mice. **(D)** Serum concentration (pg/mL) of TNF-α at day 14 only for WT versus TCF cKO-transplanted mice. **(E)** Serum concentration (pg/mL) of IFN-γ over time for WT versus TCF cKO-transplanted mice. **(F)** Serum concentration (pg/mL) of IL-2 over time for WT versus TCF cKO-transplanted mice. For **A, B**, and **D**, individual points are shown with mean and SD. For **C**, **E**, and **F**, mean and SD only is shown. All data were analyzed with one-way or two-way ANOVA or Student’s t-test, and **** means p-value ≤ 0.0001. N=3-5 per group for A-B with data from two experiments shown, N=6 per group for C-F, summary data for C,E,F and individual points in D from one representative experiment.

We also took blood from these recipient mice and obtained serum, which we tested for various cytokines using a LEGENDplex ELISA kit (Biolegend). TNF-α was found to be increased at day 7 in TCF cKO T cells, but then dropped below WT levels by day 14 (Fig. 5C). By day 14, the trend towards decreased TNF-α in serum of mice transplanted with TCF cKO donor cells was already visible, suggesting a progressive drop in TNF-α production by donor cells (Fig.5D). IFN-γ was also increased for TCF cKO T cells at day 7, and dropped to WT levels by day 14 (Fig.5E). The difference in IFN-γ results from serum and donor splenocytes suggests that loss of TCF-1 in donor cells results in increased production of IFN-γ from host cells early on, compared to in WT-transplanted mice. These data also suggest that TNF-α is more directly critical for GVHD damage than IFN-γ is. Finally, IL-2 expression in serum was increased at day 7 in TCF cKO-transplanted mice, but then dropped to WT levels by day 14 (Fig.5F). This correlated with the increased proliferation seen at day 7 with EdU, and the hypothesized drop in proliferation later on. These data suggest that allotransplanted TCF cKO T cells are more activated early on in the response, but are less active (or less present) later on.

### Loss of TCF-1 changes the suppressive profile of mature CD4 T cells

Given that loss of TCF-1 altered mature CD4 T cell phenotype and function, we sought to understand whether suppressive markers on the T cells were also altered. Expression of markers such as CTLA-4, PD-1, and PD-L1 play a role in how alloactivated T cells interact with other cells. To examine this, we allotransplanted recipient BALB/c mice with WT or TCF cKO CD4 T cells, and phenotyped donor cells from spleen and liver at day 7, day 14, and day 21 post-transplant. PD-1 expression was significantly reduced in the spleen and liver for TCF cKO CD4 T cells (Supp. Fig. 5A-B). This suggests that TCF-1-deficient cells may be less sensitive to suppression by other cells expressing PD-L1, and therefore may be more activated during disease(29). We also found that expression of PD-L1 was increased in the liver only for TCF cKO CD4 T cells at day 14 (Supp. Fig.5C-D. It is unclear what role PD-L1 may play when expressed on T cells, but it is thought to promote apoptosis of and suppress activation of other nearby T cells (30). This suggests that TCF-deficient CD4 T cells may hinder activation and survival of other T cells. Therefore, loss of TCF-1 promotes a T cell phenotype that is less susceptible to suppression yet potentially more suppressive of other cells. Finally, expression of CTLA-4 was not impacted by loss of TCF-1 (Supp. Fig.5E-F). This suggests that TCR signaling activity is not aberrantly regulated by CTLA-4 in TCF-deficient CD4 T cells, so the cells are likely not over- or understimulated through the TCR. Therefore, loss of TCF-1 changes the suppressive profile of mature CD4 T cells, but does not disturb control of TCR signaling by CTLA-4.

### TCF-1 contributes to control of cytokine and immune signaling pathways in donor CD4 T cells

To understand the molecular mechanisms behind the changes we saw in the TCF cKO donor CD4 T cells, and to understand the role of TCF-1 in regulating gene expression in these cells, we employed RNA sequencing. We allotransplanted mice with WT or TCF cKO CD3 T cells and BALB/c BM as described above. A FACS-sorted pre-transplant sample of CD4+ donor cells was taken and stored in Trizol. At day 7 post-transplant, donor T cells were sorted back from spleen and liver of recipients using H2Kb, CD3, CD4, and CD8. The sorted cells were all stored in Trizol, then sent to the SUNY Upstate Molecular Analysis Core for RNA extraction and library prep (https://www.upstate.edu/research/facilities/molecular-analysis.php). Prepped samples were then sequenced at the University of Buffalo Genomics Core (paired end sequencing on an Illumina NovaSeq 6000, http://ubnextgencore.buffalo.edu) (Fig. 6). After analyzing the data in Partek Flow, we found that for both pre- and post-transplant samples, WT and TCF cKO donor CD4 T cells clustered separately (Fig. 6A), (Supp. Fig.6A-C). In addition, WT and TCF cKO donor CD4 T cells appeared to be more similar genetically after allo-transplantation, as the number of significantly different genes was much higher in pre-transplant spleen than in post-transplant spleen and liver (Fig. 6B). In pre-transplant spleen, upregulation of Ccr2, Cxcr6, Ccr3 (chemokine receptors) and Mki67 (proliferation) was observed in TCF cKO donor cells, with downregulation of Tox2 (exhaustion) (Fig. 6C). In post-transplant spleen and liver, Ccr9, Ccl5, and Cxcr5 were altered by loss of TCF-1 in donor CD4 T cells (Fig. 6D). This suggests that chemokines and receptors are indeed regulated in mature and alloactivated CD4 T cells by TCF-1, as shown by our qPCR data.

**Figure 6:**
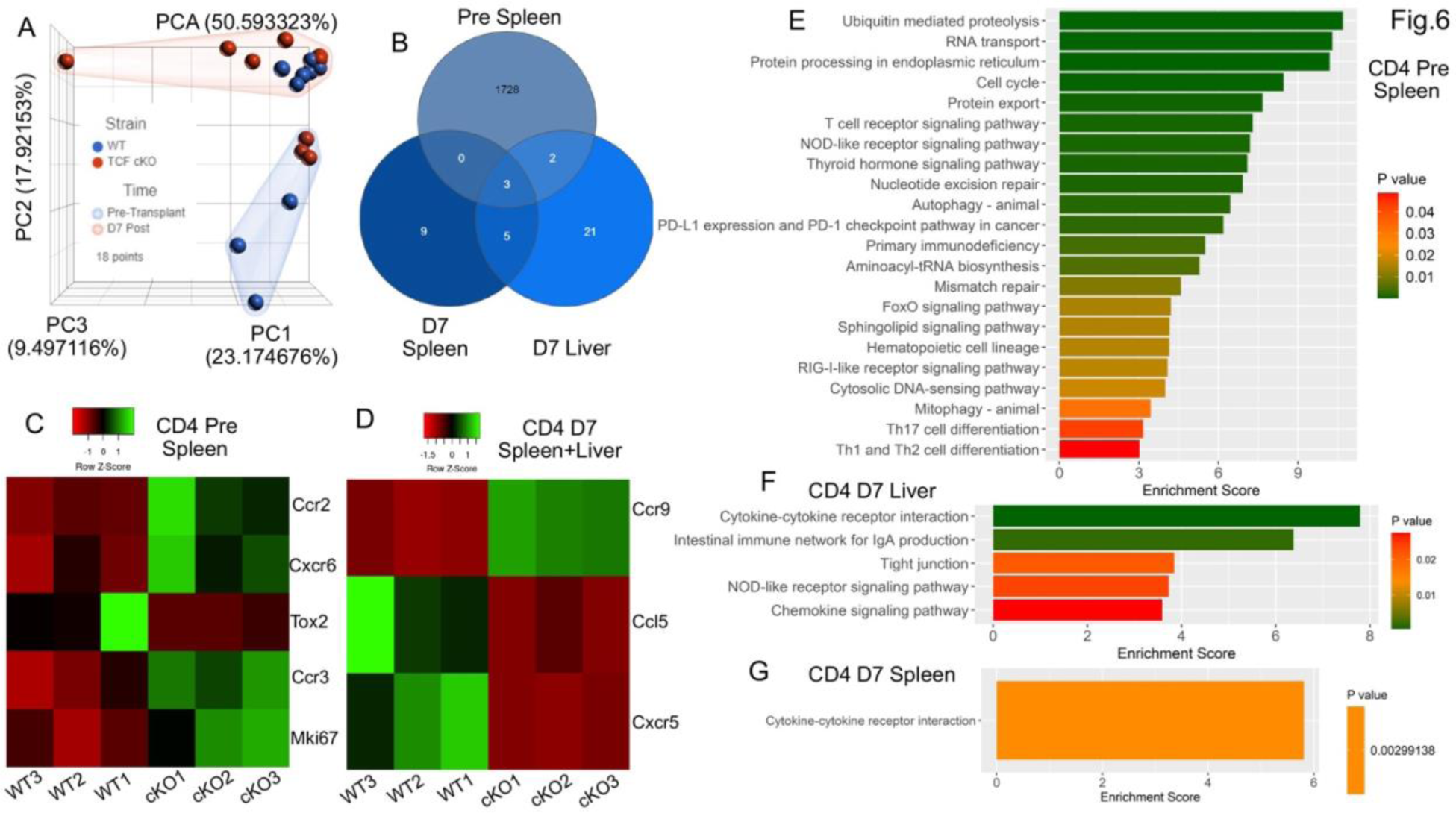
TCF-1 contributes to control of cytokine and immune signaling pathways in donor CD4 T cells. Recipient Balb/c mice were allotransplanted with Balb/c BM and 1×10^6^ WT or TCF cKO donor CD3 T cells as before. Pre-transplant donor CD4 T cells were FACS-sorted from spleens of donor mice prior to transplant, and at day 7 post-transplant, donor CD4 T cells were FACS-sorted back from recipient spleen and liver. Cells were sorted into Trizol, then RNA was extracted and paired-end sequencing was done on an Illumina sequencer. Data were analyzed using Partek Flow. **(A)** PCA analysis showing clustering of donor T cells from WT and TCF cKO mice. Points are colored by strain (blue is WT, red is TCF cKO), and highlighted by timepoint (blue grouping is pre-transplant, red grouping is d7). **(B)** Venn diagram of number of differentially expressed genes found at each time point, using criteria FDR ≤0.05 and fold change >2 or ≤ −2. **(C-D)** Heatmap showing select significantly changed genes in donor CD4 T cells from **(C)** pre-transplant spleen, or **(D)** post-transplant spleen and liver. **(E-G)** Pathway analysis for WT versus TCF cKO CD4 donor T cells at each time point and organ. Significantly altered pathways (p-value ≤ 0.05) are shown, with enrichment score plotted and p-value indicated by fill color. **(E)** Pre-transplant spleen, **(F)** Day 7 liver, and **(G)** Day 7 spleen are shown. N=3 samples per group and condition, one experiment shown (done once).

Pathway enrichment analysis showed that cell cycle control, T cell differentiation, TCR signaling, FoxO signaling, and immune signaling pathways (such as NOD and RIG-I) were altered by loss of TCF-1 in pre-transplant donor cells (Fig. 6E). Post-transplant, cytokine/cytokine receptor interaction genes were affected in both liver and spleen, and in liver only, other pathways such as intestinal immunity and NOD signaling were also impacted by loss of TCF-1 (Fig. 6F-G). Interestingly, WT and TCF cKO donor CD4 T cells appear to be more different prior to transplant as compared to post-transplant, because the number of pathways disrupted is fewer in post-transplant WT versus TCF cKO cells (Fig 6E-G). Therefore, TCF-1 controls numerous immune genes and pathways in mature and alloactivated CD4 T cells, and loss of this critical factor alters expression of these programs, ultimately affecting T cell function and phenotype.

## Discussion

T Cell Factor-1 (TCF-1) is a T cell transcription factor that is known to be critical for T cell development, activation, and in some cases, responses to pathogens (31). However, it unclear whether TCF-1 may regulate alloactivated T cells during responses to alloantigen. TCF-1 is also critical for supporting CD4 versus CD8 lineage maintenance in developing T cells (6, 32). It was also unclear whether this lineage support is maintained by TCF-1 even in mature T cells (33, 34). To address these gaps in knowledge, we studied loss of TCF-1 specifically in T cells during allo-HSCT leading to GVHD. Using a unique mouse strain with T cell-specific deletion of TCF-1 (rather than global deletion), we were able to study the role of TCF-1 in mature and alloactivated CD4 T cells (20). This manuscript, together with our companion manuscript, shows that TCF-1 does regulate mature CD4 and CD8 alloactivated T cells during GVHD (and GVL for CD8 cells). We discovered that TCF-1 appears to have a unique role in mature and alloactivated CD4+ versus CD8+ T cells, potentially due to its ability to bind different lineage-specific genes to help maintain identity in these disparate subsets.

Here, we showed that loss of TCF-1 in CD4 T cells leads to reduced severity and persistence of GVHD symptoms. Tissue damage in the target organs (liver, skin, and small intestine) was also reduced when donor cells were TCF-1-deficient. This suggests that TCF-1 is indispensable for GVHD damage, and normally promotes damage to healthy host tissues driven by donor T cells. T cell-mediated damage is a consequence of donor T cell migration, proliferation, survival, and cytokine production. If any or all of these functions are disrupted, then alloreactive T cells will be unable to induce GVHD. However, it is not ideal to fully disrupt all of these functions, because the same processes also mediate protection against infection post-transplant (35, 36). Here, we show that survival is reduced and exhaustion is increased in donor CD4 T cells from TCF-1-deficient mice. Interestingly, there appears to be a migration defect in TCF-1 cKO donor T cells as well; however, given that chemokine and chemokine receptor expression indicates that migration should be possible in these cells, it is more likely that this apparent defect stems from a reduction in proliferation or survival.

During GVHD, production of cytokines by donor cells and host tissues causes damage to nearby healthy host cells (37–39). Cytokine production was affected by loss of TCF-1, as donor T cells from TCF cKO mice showed decreased production of TNF-α, but not IFN-γ. When serum from recipient mice was tested, there was an early (day 7) increase in production of TNF-α and IFN-γ, as well as IL-2, suggesting that TCF cKO donor cells are initially highly activated. Later on (day 14 and 21), levels of all three cytokines dropped. For IFN-γ, donor cell production was not affected by loss of TCF-1, but serum production increased early on. This is likely from increased production by host cells when in the presence of TCF cKO donor cells. This suggests that despite early increased activation, cytokine production by TCF cKO donor cells quickly reduces post-transplant, allowing disease to resolve.

We also found that TCF-1-deficient donor CD4 T cells are more exhausted in the liver and trending towards more exhausted in the spleen at 7 days post-transplant than WT donor T cells. Exhausted cells are unable to respond to antigen in the proper manner, so if the donor T cells mediating GVHD damage become exhausted, they will no longer continue to cause damage(40). Donor T cells must also proliferate during GVHD to replace effector cells, which die quickly but drive damage to tissues (41). Proliferation of TCF cKO donor CD4 T cells was also initially higher in spleen and liver of transplanted mice, which correlates with the increase in serum IL-2. However, by day 14, the number of donor cells able to be sorted from recipient mice is vastly reduced for TCF cKO donors compared to WT. Therefore, we hypothesize that despite the higher early activation and proliferation of TCF cKO T cells, these donor cells rapidly become exhausted, stop proliferating, and die off. This would lead to reduced presence of donor cells in the host after day 7, which correlates well with the observed drop in symptom severity after the peak at day 7. Importantly, many of the functional changes we observed showed trends rather than statistical significance between WT and TCF cKO donor cells. We expect that this is because day 7 marks the beginning of the differences in disease outcome between groups, and the differences become much more pronounced after this time point, as supported by GVHD clinical scores and survival data. Therefore, we expect that as the disease outcomes separate more between groups, the differences in T cell function would also become even more significant. While TCF cKO donor cells may function similarly to WT cells early on, after day 7 they clearly begin to function differently, leading to changes in tissue damage and disease score. We hypothesize that after day 7, TCF cKO donor T cells become exhausted and stop proliferating and producing cytokines, allowing resolution of a usually persistent disease state (41).

To examine the mechanism behind these functional changes, we phenotyped TCF cKO and WT donor T cells prior to transplant. Despite having no effect on Eomes, T-bet, or CD122, loss of TCF-1 led to increased polarization of CD44 expression. This manifested as expansion of naive CD4 T cells, with a concomitant decrease in activating/transition CD4 T cells from the TCF cKO donor mice. This suggests that TCF-1 normally promotes activation of T cells out of the naive state for CD4 T cells, and loss of this factor allowed more cells to remain naïve (42). TCF-1 is known to be highly expressed in naive T cells (34), and naive cells drive GVHD damage (15, 43). Thus, it is clear that despite expansion of CD4 naive T cells in these TCF cKO mice, functional and genetic changes to CD4 T cells caused by loss of TCF-1 prevent exacerbation of GVHD damage. These phenotypic effects were cell-intrinsic, indicating a direct role of TCF-1 in controlling mature CD4 T cell phenotype. Thus, TCF-1 directly and cell-intrinsically produces changes in CD4 T cell phenotype, which can impact alloactivated T cell functioning. We also discovered that loss of TCF-1 altered expression of PD-1 and PD-L1 by CD4 T cells following allotransplant. This suggests that the donor T cell suppressive profile is altered following transplant. No changes were observed for CTLA-4, suggesting that TCR signaling regulation is not disrupted by loss of TCF-1.

Finally, to examine what changes occurred to the genetic program following loss of TCF-1, we performed RNA sequencing on pre- and post-transplant donor cells as an unbiased approach. Gene expression by T cells changes drastically during alloactivation (44), and we sought to understand how mature T cells lacking TCF-1 were different from those with TCF-1, both before and after alloactivation. We FACS-sorted splenic donor CD4 T cells pre-transplant, and both splenic and liver-derived donor cells post-transplanted (from recipients). We found that many immune-related genes and pathways were altered by loss of TCF-1 in donor cells, both pre- and post-transplant. Interestingly, allotransplantation appears to induce genetic programs that are similar in both WT and TCF cKO donor cells, because the number of altered pathways and genes when comparing strains was less for post-transplant cells than for pre-transplant cells. This means that pre-transplant donor cells are more different based on expression of TCF-1 than post-transplant cells are. Of note, the most significantly impacted pathway in post-transplant cells (from spleen or liver) was cytokine/cytokine receptor interactions. This highlights the importance of our finding that TCF cKO donor cells trend towards less TNF-α production, suggesting that TNF-α is critical for driving GVHD damage (45), and therefore helping to explain our observation that TCF cKO-transplanted mice have less severe GVHD. We also found that intestinal immunity and NOD signaling (46) were altered post-transplant, again helping to explain the reduced gut symptoms seen in TCF cKO-transplanted mice (47). Alterations in the chemokine signaling pathway were also detected, supporting our qPCR data showing that TCF-1 controls chemokine receptor expression on mature CD4 T cells (48, 49). Pre-transplant, pathways such as cell cycle, T cell differentiation/signaling, and immune signaling pathways were affected by loss of TCF-1, helping to explain the altered phenotype and function of mature TCF cKO CD4 T cells (50). Overall, this sequencing data shows that TCF-1 is not only important in T cell development, but is also critical for regulation of mature and alloactivated T cells as well.

Our companion manuscript addresses the role of TCF-1 in mature alloactivated CD8 T cells in this model. Interestingly, the effects on phenotype, gene expression changes, and chemokine/chemokine receptor expression were different for CD8 versus CD4 T cells. The ability of TCF-1 to maintain specific lineage traits for CD4 and CD8 T cells seems to be maintained in mature alloactivated T cells as well, which was previously unknown. Overall, our data show that TCF-1 directly regulates alloactivated mature CD4 T cells, which was previously unknown. They also show that CD4 versus CD8 mature alloactivated T cells are both regulated in different ways by TCF-1, leading to disparate effects of TCF-1 deficiency in these cells. However, loss of TCF-1 in either cell type does produce a clinically optimal phenotype, with reduced host tissue damage and GVHD severity over time. Thus, modulation of TCF-1 or critical downstream factors may prove beneficial for reducing GVHD severity following allo-HSCT. Additionally, TCF-1 modulation in T cells may be useful in other T cell-mediated disorders.

## Materials and Methods

### Mouse Models

For transplant experiments, the following female donor mice were used: B6-Ly5.1 (CD45.1+, “WT” or B6.SJL-Ptprca Pepcb /BoyCrl, 494 from Charles River), C57Bl/6j (CD45.2+, “WT”,”, 000664 from Jackson Laboratories), Tcf7 flox x CD4cre (referred to here as “TCF cKO”, obtained from Dr. Jyoti Misra Sen at at NIH by permission of Dr. Howard Xue, and bred in-house (20), or CD4cre (022071 from Jackson Laboratories, bred in-house). These donors were age-matched to each other and to recipients as closely as possible. Recipient mice for transplant experiments were female BALB/c mice (CR:028 from Charles River, age 8 weeks or older). Recipient mice for chimera experiments were Thy1.1 mice (B6.PL-Thy1a/CyJ, 000406 from Jackson Labs).

### Bone Marrow Transplants

Short-term and long-term bone marrow transplant models were employed. BALB/c recipient mice were irradiated twice with 400 cGy of x-rays (total dose 800 cGy), with a rest period of at least 12 hours between doses, and 4 hours of rest prior to transplantation. T cells (total CD3+ or CD4+) were separated from WT, CD4cre, or TCF cKO spleens using CD90.2 or CD4 microbeads and LS columns (Miltenyi, CD4: 130-117-043, CD90.2: 130-121-278, LS: 130-042-401). The cells were then injected IV into the tail vein in PBS. The recipient mice received 1×10^6^ T cells per mouse, along with 10×10^6^ bone marrow cells collected from BALB/c mice. Bone marrow was T-cell depleted with CD90.2 MACS beads (130-121-278 from Miltenyi) and LD columns (130-042-901 from Miltenyi). At day 7, the recipients were humanely euthanized, and serum, spleen, lymph nodes, small intestine, skin, or liver was collected, depending on the experiment. For migration experiments, the donor cells were a 1:1 ratio of WT (CD45.1):WT(CD45.2), or WT:TCF cKO CD3 T cells (total 1×10^6^ cells). For GVHD experiments, the recipient mice were weighed and given a GVHD score from day 1 to day 60, and group survival was analyzed. For mixed bone marrow chimera experiments, a 1:4 ratio of WT (CD45.1) to TCF cKO (CD45.2) bone marrow (total 50×10^6^ cells) was injected into Thy1.1 mice, and the mice were rested for 9 weeks. At 9 weeks post-transplant, tail vein blood was collected and stained with anti-CD45.1 and anti-CD45.2 to detect the two donor cell types. At 10 weeks, these chimeras were euthanized and their spleens were processed and stained for phenotyping by flow cytometry.

### Flow Cytometry, Sorting, and Phenotyping

For phenotyping experiments and pre-transplant sorted cells, splenocytes were obtained from WT and TCF cKO mice. For all other experiments, cells were obtained from transplanted recipients. Cells were incubated with RBC Lysis Buffer (00-4333-57 from eBioscience) to remove red blood cells when necessary. After processing, cells were stained in MACS buffer (1x PBS with EDTA and 4g/L BSA) with extracellular markers, and were incubated for 30 minutes on ice. Cells were then washed and run on a BD LSRFortessa flow cytometer to collect data. If intracellular markers were used, cells were washed after extracellular staining, then fixed overnight using buffers from the Fix/Perm Concentrate and Fixation Diluent from FOXP3 Transcription Factor Staining Buffer Set (eBioscience cat. No. 00-5523-00). The following day, cells were washed in Perm buffer from the same kit, and were stained with intracellular markers in Perm buffer for 40 minutes at room temperature. Stained cells were resuspended in FACS buffer (eBioscience cat. No. 00-4222-26) and transferred to flow tubes. All antibodies were used at 1:100 dilution, and were purchased from Biolegend or eBioscience. The cells were then washed and run on a BD LSRFortessa. For cell sorting, cells were stained in the same manner and run on a BD FACSAria, equipped with cold-sorting blocks. Cells were sorted into sorting media (50% FBS in RPMI) for maximum viability, or Trizol for RNAseq/qPCR experiments. All flow cytometry data was analyzed using FlowJo software v9 (Treestar). Depending on the experiment, antibodies used were: anti-CD4 (FITC, PE, BV785, BV21), anti-CD8 (FITC, PE, APC, PerCP, Pacific Blue, PE/Cy7), anti-CD3 (BV605 or APC/Cy7), anti-H2Kb-Pacific Blue, anti-H2Kd-PE/Cy7, anti-CD122 (FITC or APC), anti-CD44 (APC or Pacific Blue), anti-CD62L (APC/Cy7), anti-TNF-α-FITC, anti-IFN-γ-APC, anti-Eomes (AF488 or PE/Cy7), anti-T-bet-BV421, anti-CD45.2-PE/Dazzle594, anti-CD45.1-APC, anti-Ki67 (PE or BV421), anti-PD-1-BV785, anti-CTLA-4-PE, and anti-PD-L1 (PE or PE/Dazzle594), anti-Annexin V-FITC, LIVE/DEAD Near IR, anti-EdU-AF647 (click chemistry kit), and anti-TOX-APC

### Histology

At day 7, day 14, and day 21 post-transplant, organs were removed from WT or TCF cKO CD4 T cell-transplanted recipient mice. The spleen, liver, small intestine, and skin from back and ear were removed and fixed in 10% neutral buffered formalin. Tissue pieces were sectioned and stained with hemotoxylin and eosin by the Histology Core at Cornell University (https://www.vet.cornell.edu/animal-health-diagnostic-center/laboratories/anatomic-pathology/services). Stained slides were then imaged and given a GVHD grade by a pathologist at SUNY Upstate (A.M.) who was blinded to study conditions and slide identity. GVHD grade was determined on a scale of 0-4 by the following criteria: for skin - basal cell vacuolation up to frank epidermal denudation, for small intestine - single cell necrosis up to diffuse mucosal denudation, and for liver - percent of bile duct showing epithelial damage (I ≤25%, II 25-49%, III 50-74%, IV 75-100%). Links for grading criteria: http://surgpathcriteria.stanford.edu/transplant/skinacutegvhd/printable.html, http://surgpathcriteria.stanford.edu/transplant/giacutegvhd/printable.html, http://surgpathcriteria.stanford.edu/transplant/livergvhd/printable.html.

### Isolation of lymphocytes from liver, small intestine, or lymph nodes

To isolate lymphocytes from lymph nodes, the inguinal, brachial, and axillary lymph nodes were removed from euthanized mice, and ground between glass slides to create a cell suspension. Ground cells were rinsed into tubes using PBS and centrifuged to remove debris. To isolate lymphocytes from liver, livers were perfused with 5-10mL of ice cold PBS to remove RBCs before organ removal. Livers were then ground through a 70uM filter with PBS, centrifuged to remove debris, and lymphocytes were isolated by a 22 minute spin in 40% Percoll in RPMI/PBS (22 degrees C, 2200rpm, no brake, no acceleration). Isolated lymphocyte pellets were washed, lysed to remove remaining RBCs, and resuspended with PBS or MACS buffer (BSA in PBS). To isolate lymphocytes from small intestine, the intestine was removed and put in ice cold MACS buffer, opened lengthwise, washed with MACS, and epithelial cells were stripped off by a 30 min shaking incubation (37 C) in strip buffer (1x PBS, FBS, EDTA 0.5M, and DTT 1M). Guts were then cut into small pieces and digested by a 30 min shaking incubation (37 C) in digestion buffer (collagenase, DNAse, and RPMI). The tubes were then vortexed, and liquid and solid gut pieces were filtered through a 70uM filter to obtain a cell suspension. Percoll was then used to isolate lymphocytes as for liver, with no RBC lysis afterwards. Gut cells were then placed in MACS buffer for further use.

### qPCR Analysis

Mice were short-term transplanted as described above (1×10^6^ CD3 donor T cells and 10×10^6^ BALB/c BM), and at day 7, recipient mice were humanely euthanized. Splenocytes were obtained from pre- and post-transplanted mice, and FACS sorted as described above to obtain donor cells. These cells were all sorted into Trizol, then RNA was extracted using chloroform phase separation protocols (https://www.nationwidechildrens.org/Document/Get/93327). The extracted RNA was eluted using the Qiagen RNEasy Minikit (74104 from Qiagen) and run on a spectrophotometer to determine concentration. RNA was then converted to cDNA with the Invitrogen SuperScript IV First Strand Synthesis System kit (18091050 from Invitrogen) and run on a spectrophotometer to determine concentration. Master cocktail including 10ng/uL cDNA and Taqman Fast Advanced Master Mix (4444557 from Invitrogen) was prepared for each sample, and 20uL was added to each well of a 96 well TaqMan Array plate with chemokine/chemokine receptor primers (ThermoFisher, Mouse Chemokines & Receptors Array plate, 4391524). The plates were run on a Quantstudio 3 thermocycler according to manufacturer instructions for the TaqMan assay, and data were analyzed using the Design and Analysis software v2.4 (provided by ThermoFisher). Five separate recipient mice were sorted and cells were combined to make one sample for qPCR testing per condition/organ.

### Migration Assay

BALB/c mice were allotransplanted with 10×10^6^ BALB/c BM, and a 1:1 mixture of WT CD45.1+ CD3 T cells with either WT CD45.2+ cells or TCF cKO CD45.2+ cells (total 1×10^6^ cells). This 1:1 mixture was checked prior to transplant to ensure a 1:1 ratio of CD4 to CD8 T cells per strain, as well as a 1:1 ratio of each donor type (WT:WT or WT:TCF cKO). At day 7 post-transplant, recipient mice were euthanized and lymphocytes were obtained as described above from spleen, lymph nodes, liver, and small intestine. The lymphocytes were stained for CD3, CD4, CD8, and H2Kb to identify donor cells, as well as CD45.2 and CD45.1 to identify cells of each donor strain. The ratio of CD45.1 to CD45.2 cells (i.e. WT to WT or WT to TCF cKO) was calculated from flow cytometry data and analyzed.

### Cytokine Restimulation

Mice were short-term transplanted as described above (1.5×10^6^ CD3) donor T cells and 10×10^6^ BALB/c BM), and at day 7 post-transplant, recipient mice were humanely euthanized and splenocytes were obtained. The splenocytes were cultured for 6 hours at 37C and 7% CO2 with GolgiPlug (1:1000) and PBS or anti-CD3 (1ug/mL)/anti-CD28 (2ug/mL) to restimulate them. After 6 hours, the cells were removed from culture, stained for surface markers, fixed and permeabilized, then stained for the cytokines IFN-y and TNF-a using the BD Cytokine Staining kit (555028), and run on a flow cytometer.

### LEGENDplex Serum ELISA Assay

In addition to splenocytes, serum was collected from recipient mice in the cytokine experiment, and analyzed using the Biolegend LEGENDplex Assay Mouse Th Cytokine Panel kit (741043). This kit quantifies serum concentrations of: IL-2 (T cell proliferation), IFN-y and TNF-a (Th1 cells, inflammatory), IL-4, IL-5, and IL-13 (Th2 cells), IL-10 (Treg cells, suppressive), IL-17A/F (Th17 cells), IL-21 (Tfh cells), IL-22 (Th22 cells), IL-6 (acute/chronic inflammation/T cell survival factor), and IL-9 (Th2, Th17, iTreg, Th9 – skin/allergic/intestinal inflammation).

### Proliferation Assay

Mice were short-term transplanted as described above (1×10^6^ CD3 donor T cells and 10×10^6^ BALB/c BM), and recipient mice were injected at day 5-6 with 25mg/kg EdU (20518 from Cayman Chemicals) in PBS. At day 7, the recipient mice were humanely euthanized, and lymphocytes from spleen and liver were obtained. These cells were processed and stained using an EdU click chemistry kit (C10424 from Invitrogen), stained for H2Kb, CD3, CD4, and CD8, and run on a flow cytometer. In vitro culture attempts mentioned as empirical observations were done at 37 C, 7% CO2, with anti-CD3 (1ug/mL), anti-CD28 (2ug/mL), and in some cases, IL-2 (1000IU/mL).

### Death Assay

Mice were short-term transplanted as described above (1×10^6^ CD3 donor T cells and 10×10^6^ BALB/c BM), and at day 7 post-transplant, cells from spleen and liver were stained with Annexin V-FITC (V13242 from Invitrogen) and LIVE/DEAD Near IR (L34976 from Invitrogen). Annexin V and NIR were used to identify dead (Ann.V+NIR+), live (Ann.V-NIR-), and apoptotic (Ann.V+NIR-) cells. Donor T cells were identified by H2Kb, CD3, CD4, and CD8.

### Exhaustion/Activation Assay

Mice were short-term transplanted as described above (1×10^6^ CD3 donor T cells and 10×10^6^ BALB/c BM), and at day 7, lymphocytes were obtained from spleens and livers of transplanted mice. The cells were stained with anti-TOX and anti-Ki67. Donor T cells were identified by H2Kb+, CD3+, CD4+, and CD8+.

### Time course Assay

Mice were short-term transplanted as described above (1×10^6^ CD4 donor T cells and 10×10^6^ BALB/c BM), and at day 7, day 14, and day 21, lymphocytes were obtained from spleens and livers of transplanted mice. The cells were stained with anti-PD-1, anti-PD-L1, and anti-CTLA4. Donor T cells were identified by H2Kb+, CD3+, CD4+, and CD8+.

### cDNA Extraction and PCR

Donor mice were genotyped using PCR. Ear punches were taken from each mouse at 4 weeks of age, DNA was extracted, and run in a PCR reaction using the Accustart II Mouse Genotyping kit (95135-500 from Quanta Biosciences). Standard PCR reaction conditions and primer sequences from Jackson Laboratories were used for Eomes, T-bet, and CD4cre. For Tcf7, primer sequences and reaction conditions were obtained from Dr. Jyoti Misra Sen of NIH.

### RNA Sequencing

Mice were short-term transplanted as described above (1×10^6^ CD3 donor T cells and 10×10^6^ BALB/c BM), and at day 7, recipient mice were humanely euthanized. Splenocytes were obtained from pre- and post-transplanted mice, and liver cells were obtained from post-transplant mice only. For TCF cKO-transplanted mice, cells were also sorted from recipients on day 14. Cells were FACS sorted as described above to obtain donor cells. These cells were all sorted into Trizol and brought to the Molecular Analysis Core at SUNY Upstate (https://www.upstate.edu/research/facilities/molecular-analysis.php) for RNA extraction and library prep, followed by RNA sequencing analysis at the University at Buffalo Genomics Core (http://ubnextgencore.buffalo.edu). Data were provided and were analyzed using Partek Flow software. This data will be deposited at (https://www.ncbi.nlm.nih.gov/geo/).

### Statistical Analysis

All numerical data is reported as means with standard deviation unless otherwise noted. Data was analyzed for significance with GraphPad Prism v7 or v9. Differences were determined using one-way or two-way ANOVA and Tukey’s multiple comparisons tests for three or more groups, or with a student’s t-test when only two groups were used. Kaplan-Meier survival analysis was used for survival experiments. Chi-square analysis was used for GVHD histology grades. All tests were two-sided. P-values less than or equal to 0.05 were considered significant. All transplant experiments were done with N=3-5 mice per group, and repeated at least twice. Ex vivo experiments were done two to three times unless otherwise noted with at least three replicates per condition each time. RNAseq was done once with three replicates per group. qPCR was done once with one sample per condition, and 5 mice combined to make the one sample. Sorting was done once for each of these two experiments, and data were recorded for Supp. Fig. 4. The time course assay was done once with 4-6 samples per group. The chimera experiment was done once with 5 chimeras (5 samples per donor type). Data in figures are presented as mean and SD unless otherwise noted.

### Study Approval

All animal studies were reviewed and approved by the IACUC at SUNY Upstate Medical University. All procedures and experiments were performed according to these approved protocols.

## Author Contributions

RH designed and conducted experiments, analyzed data, and wrote the manuscript. MK assisted with scientific/technical research design, and edited the manuscript. MM assisted with data collection. QY provided technical and scientific advice and assisted with data analysis. AM performed blinded histology analyses. JMS provided TCF cKO mice with permission of Howard Xue and helped edit the manuscript. IF produced gene expression heatmaps for RNAseq data.

## Acknowledgments

We thank all members of the Karimi lab for helpful discussions. This research was funded in part by a grant from the National Blood Foundation Scholar Award to MK, the National Institutes of Health (NIH LRP #L6 MD0010106 and K22 (AI130182) to MK), and an Upstate Medical University Cancer Center grant (1146249-1-75632) to MK. JMS was supported by the Intramural Research Program of the National Institute of Aging We thank Dr. Howard Xue for permission to use TCF cKO mice. Flow cytometers and sorters were provided and maintained by Lisa Phelps of the SUNY Upstate Flow Core. RNA extraction and prep was done by Karen Gentile of the SUNY Upstate Molecular Analysis core, and RNA sequencing was done by the University at Buffalo Genomics Core.

---

**Summary Figure:**
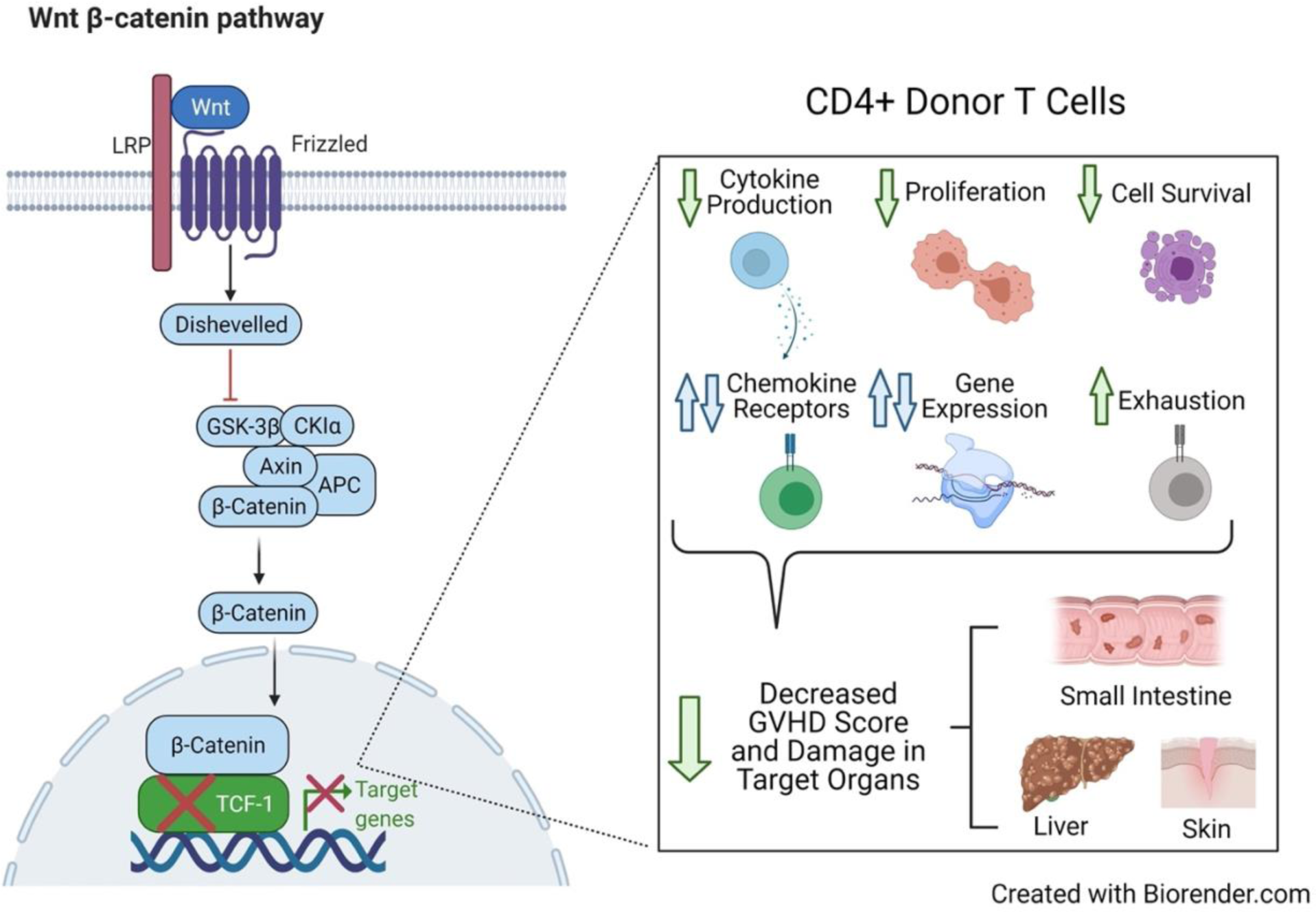
TCF-1 regulates GVHD severity and persistence by controlling mature alloactivated CD4 T cell functions and gene expression. T Cell Factor-1 (TCF-1) is a Wnt pathway transcription factor found in T cells. Loss of this factor specifically in T cells reduces GVHD severity and persistence due to changes in T cell function, including: decreased cytokine production, proliferation, and cell survival; increased exhaustion; and altered gene expression and expression of chemokine receptors. These changes lead to reduced GVHD score and less damage to target organs, including small intestine, liver, and skin.

## Supplemental Figure Legends

**Supplemental Figure 1:**
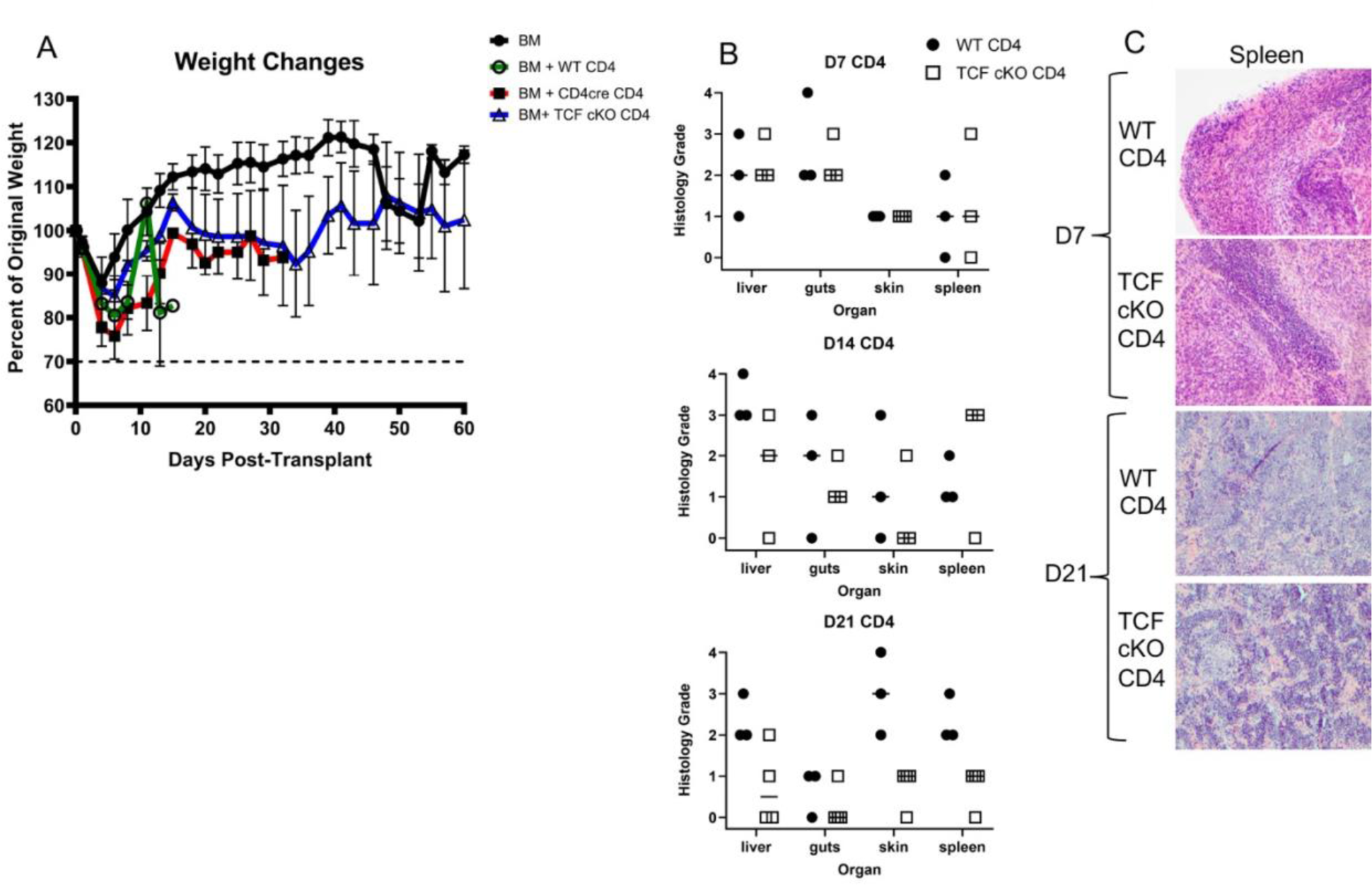
GVHD symptoms and organ damage are reduced by loss of TCF-1, related to Figure 1. Recipient mice were allotransplanted with BALB/c BM and WT, CD4cre, or TCF cKO CD4 T cells as described in Fig. 1. **(A)** Weight loss for each group of mice over the 60 days of the experiment. **(B)** Skin, spleen, liver, and small intestine were taken from allotransplanted recipient mice at d7, d14, and d21 post-transplant. A blinded pathologist (A.M.) scored each tissue section for GVHD damage (as in Fig. 1), and grades are plotted here for each section, with median indicated. **(C)** Spleen tissue sections from d7 and d21 are shown here. N=3-5 per group for A with one representative experiment shown, N=3-4 per group for B-C, one representative photo shown in **C** and individual points in **B**.

**Supplemental Figure 2:**
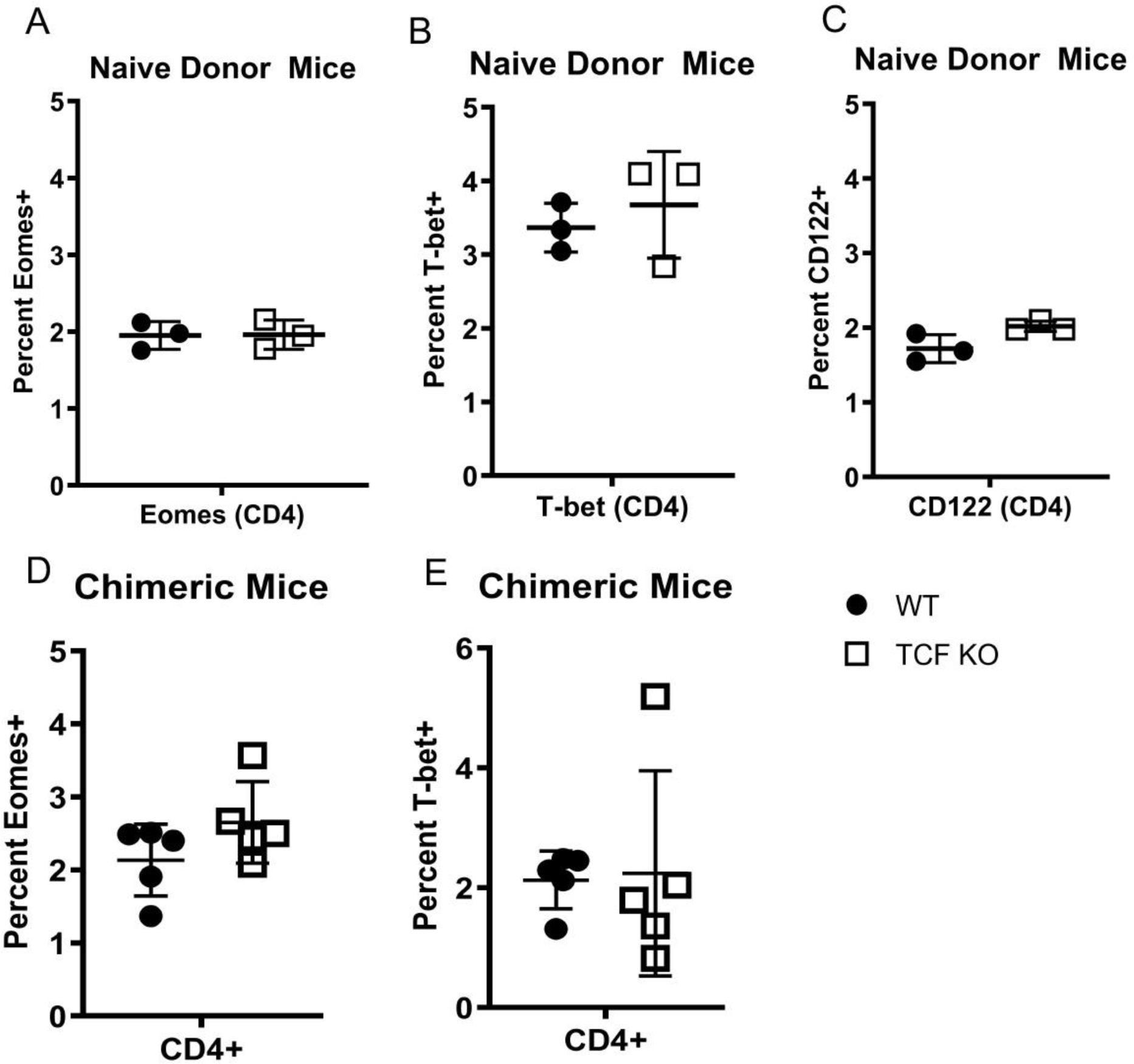
Loss of TCF-1 does not disrupt some phenotypic markers, related to Figure 2. **(A-C)** Splenocytes were taken from naive WT or TCF cKO donor mice and stained for flow cytometry. **(A)** Percent of CD4 T cells expressing Eomes. **(B)** Percent of CD4 T cells expressing T-bet. **(C)** Percent of CD4 T cells expressing CD122. **(D-E)** Thy1.1 mice were lethally irradiated and reconstituted with a 1:4 (WT:TCF cKO) mixture of bone marrow cells. Blood was checked by flow cytometry at 9 weeks to ensure reconstitution, and splenocytes were taken at 10 weeks for flow cytometry phenotyping. WT donor cells were identified by CD45.1, while TCF cKO donor cells were identified by CD45.2 **(D)** Percent of chimeric CD4 T cells expressing Eomes. **(E)** Percent of chimeric CD4 T cells expressing T-bet. N=3-5 per group for **A-C** with one representative experiment shown, N=5 per group for D-E, one experiment shown (done once).

**Supplemental Figure 3:**
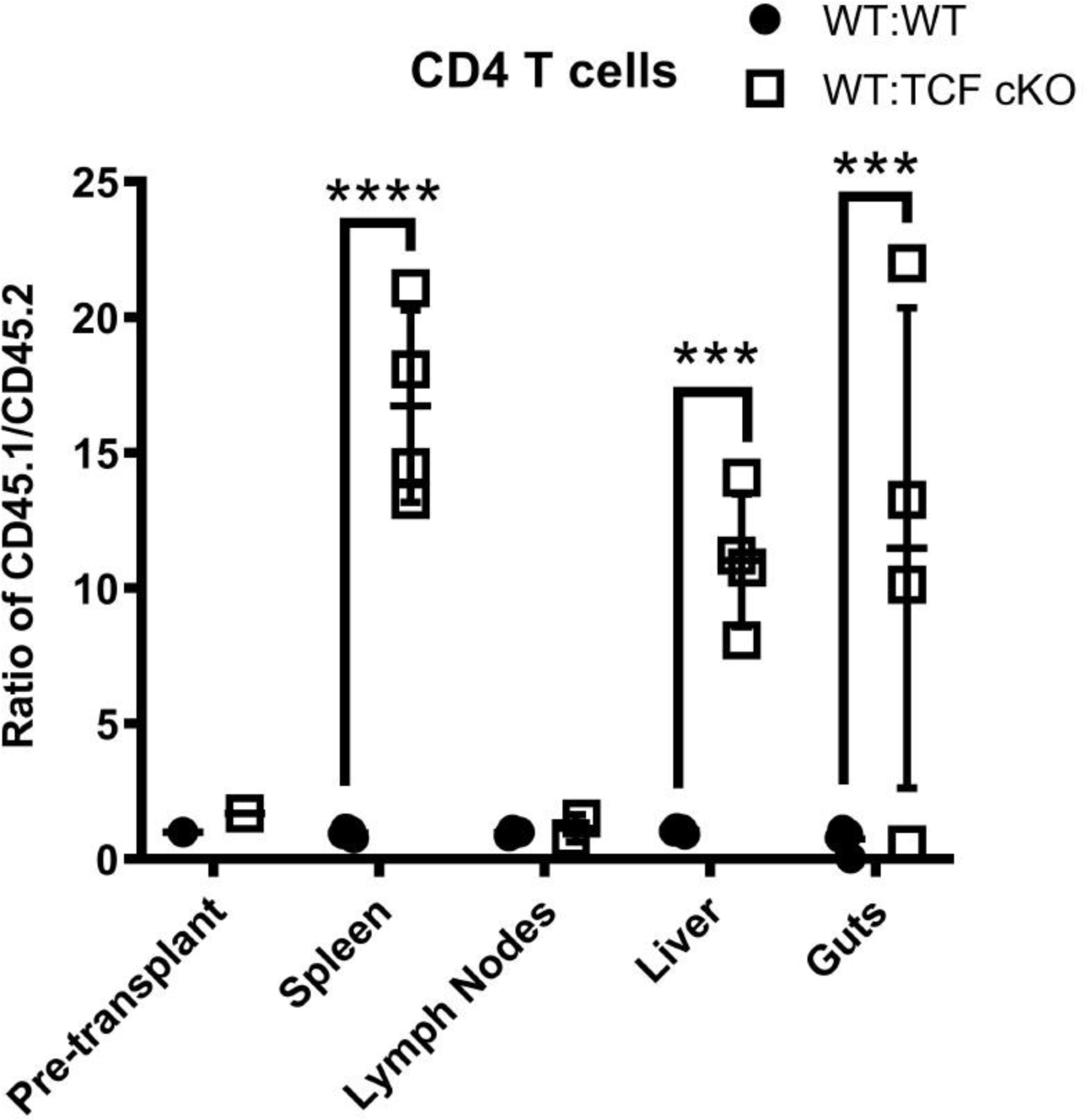
Loss of TCF-1 alters migration of T cells, related to Figure 3. Recipient BALB/c mice were lethally irradiated and allotransplanted with 10×10^6^ BALB/c bone marrow (BM) cells and 1×10^6^ CD3 T cells. The T cells were mixed at a 1:1 ratio of CD3 T cells from WT (CD45.1) and TCF cKO (CD45.2) donor mice, or WT (CD45.1) and WT (CD45.2) control mice. Pre-transplant and at day 7 post-transplant, the ratio of WT:TCF cKO (or WT:WT) cells was assessed in spleen, lymph nodes, liver, and small intestine (“guts”) using flow cytometry. The ratio of cell percentages in each organ was calculated as the ratio of WT (CD45.1) to TCF cKO or WT (CD45.2). Individual points are shown with mean and SD, data were analyzed with two-way ANOVA, **** means p-value ≤ 0.0001, and *** means p-value ≤ 0.001. N=3-4 per group with one representative experiment shown, except pre-transplant (n=1 each).

**Supplemental Figure 4:**
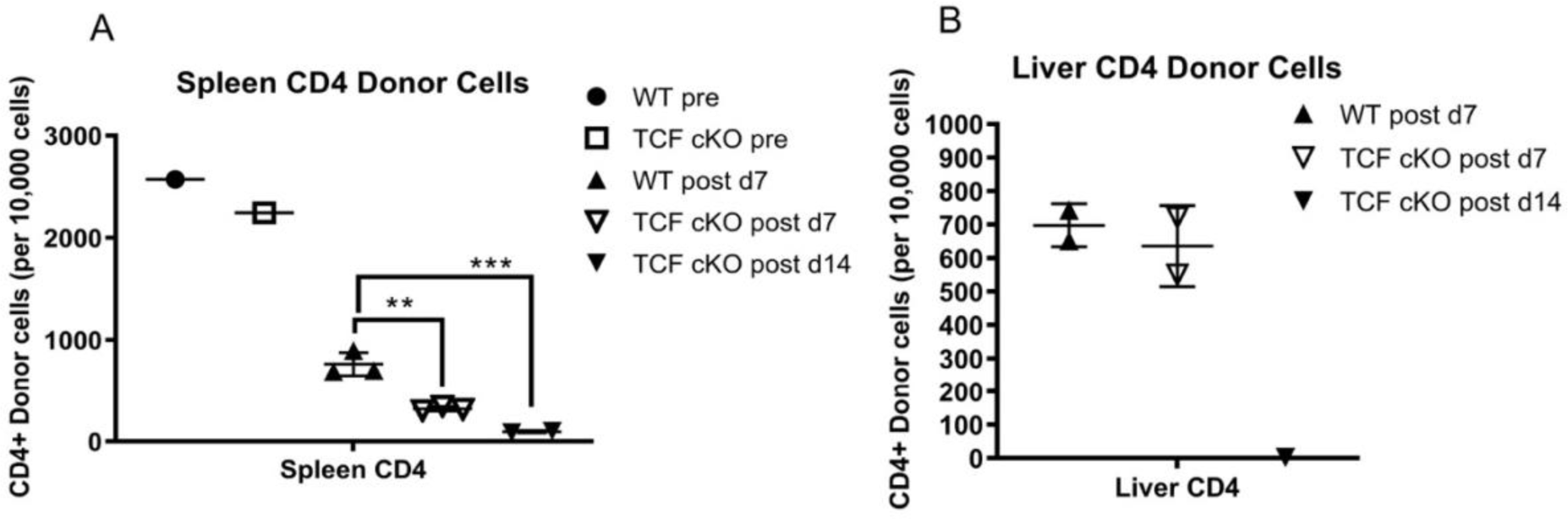
Donor cell numbers are reduced when TCF-1 is lost, related to Figure 4. Recipient mice were allotransplanted with BALB/c BM and WT or TCF cKO CD3 T cells. This was done for the RNA sequencing and qPCR experiments described, where donor cells must be FACS-sorted back from recipients post-transplant. Number of pre- and post-transplant CD4+ donor T cells obtained by sorting from donor or recipient mice (using antibodies for H2Kb, H2Kd, CD3, CD4, and CD8) is shown per 10,000 sorted cells. Number of cells for each donor type and timepoint in (A) spleen and (B) liver, with individual points and mean with SD plotted. Data were analyzed with two-way ANOVA, ** means p-value ≤ 0.01, and *** means p-value ≤ 0.001. N=1-3 per group for A (1 pre, 2 d14, 3 d7), n=2 for B, one experiment shown (done once).

**Supplemental Figure 5:**
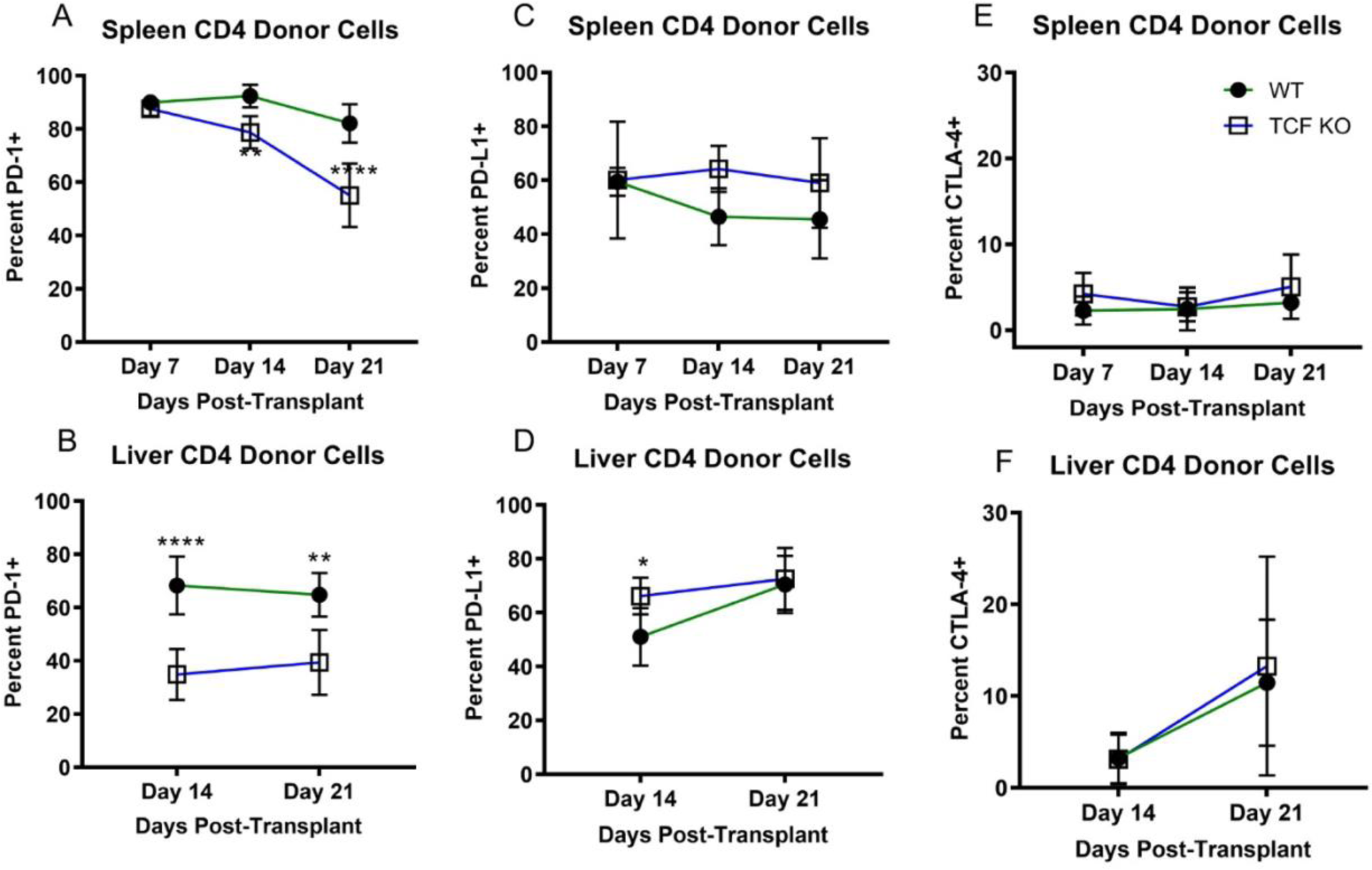
Loss of TCF-1 changes the suppressive profile of mature CD4 T cells, related to Figure 5. Recipient mice were allotransplanted with BALB/c BM and WT or TCF cKO CD4 donor T cells, as described before. At day 7, day 14, and day 21 post-transplant, spleens and livers (d14 and d21 only) were obtained from euthanized recipient mice. Lymphocytes were obtained and stained for H2Kb, CD3, CD4, and CD8, as well as PD-1, PD-L1, and CTLA-4. **(A)** Expression of PD-1 on CD4+ donor cells in spleen over time. **(B)** Expression of PD-1 on CD4+ donor cells in liver over time. **(C)** Expression of PD-L1 on CD4+ donor cells in spleen over time**. (D)** Expression of PD-L1 on CD4+ donor cells in liver over time. **(E)** Expression of CTLA-4 on CD4+ donor cells in spleen over time. **(F)** Expression of CTLA-4 on CD4+ donor cells in liver over time. All graphs show mean and SD plotted. Data were analyzed with two-way ANOVA, * means p-value ≤ 0.05, ** means p-value ≤ 0.01, and **** means p-value ≤ 0.0001. N=3-5 per group with summary data for one experiment shown (done once).

**Supplemental Figure 6:**
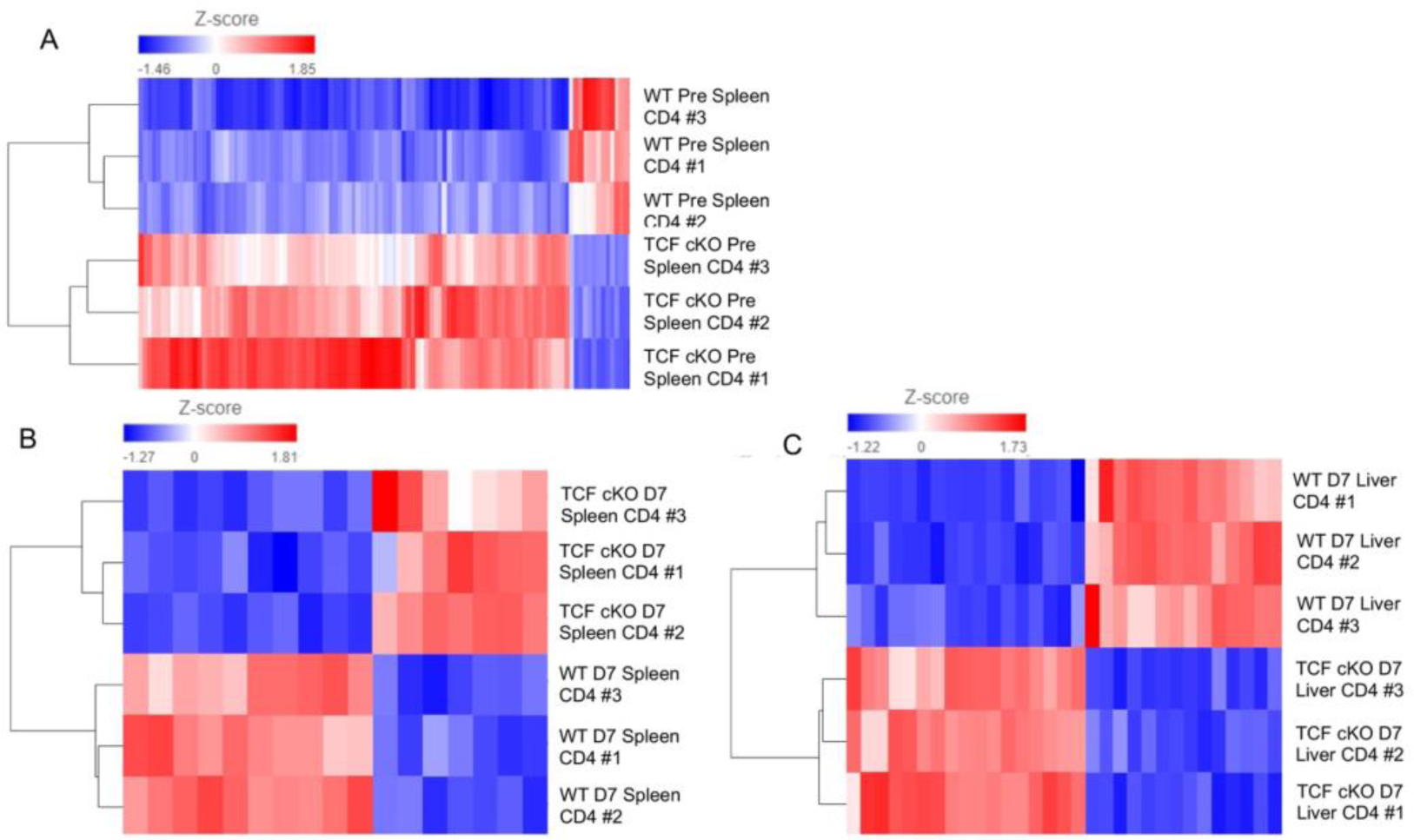
Donor cell samples cluster tightly based on absence or presence of TCF-1, related to Figure 6. As in Fig. 6, recipient BALB/c mice were allotransplanted with BALB/c BM and WT or TCF cKO donor CD3 T cells. Pre-transplant donor CD4 T cells and day 7 post-transplant donor CD4 T cells were FACS-sorted from donor or recipient mice, respectively. Cells were sorted into Trizol, RNA was extracted, paired-end sequencing was done, and data were analyzed in Partek Flow. Hierarchecal clustering was performed for **(A)** pre-transplant spleen samples, **(B)** D7 spleen samples, and **(C)** D7 liver samples. Z-score scale used for heatmap is shown above each map. N=3 per group and condition, one experiment shown (done once).

